# Mitochondrial Fatty Acid Synthesis and Mecr Regulate CD4^+^ T Cell Function and Oxidative Metabolism

**DOI:** 10.1101/2024.07.08.602554

**Authors:** KayLee K. Steiner, Arissa C. Young, Andrew R. Patterson, Erin Q. Jennings, Channing Chi, Zaid Hatem, Darren R. Heintzman, Ayaka Sugiura, Emily N. Arner, Allison E. Sewell, Matthew Z. Madden, Richmond Okparaugo, Emilia Fallman, Katherine N. Gibson-Corley, Kelsey Voss, Denis A. Mogilenko, Jeffrey C. Rathmell

## Abstract

**Summary:** We show that the mitochondrial fatty acid synthesis gene *Mecr* shapes CD4^+^ T cell metabolism and function. It may be targeted in inflammatory diseases and provides rationale to consider the immunological state of patients with mitochondrial disease.

Lipid metabolism is fundamental to CD4^+^ T cell metabolism yet remains poorly understood across subsets. Therefore, we performed targeted *in vivo* CRISPR/Cas9 screens to identify lipid-associated genes essential for T cell subset functions. These screens established mitochondrial fatty acid synthesis (mtFAS) genes *Mecr, Mcat* and *Oxsm* as highly impactful. Of these, the inborn error of metabolism gene *Mecr* was most dynamically regulated. Effector and memory T cells were reduced in *Mecr*^fl/fl^; *Cd4*^cre^ mice, and MECR was required for activated CD4^+^ T cells to efficiently proliferate, differentiate, and survive. *Mecr*-deficient T cells also had decreased mitochondrial respiration, reduced TCA intermediates, and accumulated intracellular iron, which contributed to cell death and sensitivity to ferroptosis. Importantly, *Mecr*-deficient T cells exhibited fitness disadvantages in inflammatory, tumor, and infection models. mtFAS and MECR thus play important roles in activated T cells and may provide targets to modulate immune functions in inflammatory diseases. The immunological state of MECR- and mtFAS-deficient patients may also be compromised.

## Introduction

T cell metabolism is fundamentally integrated with cell differentiation and function, as each T cell subset requires a distinct metabolic program to reach its potential (Michalek et al., 2011; Wilfahrt and Delgoffe, 2024). T cell metabolism can therefore be targeted to affect the outcome of a variety of inflammatory diseases and cancer (Lim et al., 2022; Wilfahrt and Delgoffe, 2024). Lipid metabolism controls a range of T cell biosynthetic, signaling, and energetic processes but differentially affects T cell subsets. Th1, Th17 CD4^+^ and effector CD8^+^ T cells increase lipid synthesis upon activation while T regulatory (Treg) cells rely on fatty acid ß-oxidation (Field et al., 2020; Kanno et al., 2023; Michalek et al., 2011). These pathways can modulate T cell fate and inflammatory disease models. Inhibiting lipid metabolic protein acetyl-CoA carboxylase 1 (ACC1) *in vivo* dysregulates the formation of Th17 CD4 T cells, promotes Treg cell development, and prevents Th17-mediated autoimmune disease EAE (Berod et al., 2014; Kao et al., 2023). Conversely, the uptake of oxidized lipids can contribute to T cell dysfunction in tumors (Xu et al., 2021). The roles of specific lipid metabolic enzymes or processes in inflammatory diseases, however, have not been explored through unbiased genetic screens.

Mitochondrial fatty acid synthesis (mtFAS) generates mitochondrial lipids independent of the cytosolic fatty acid synthesis (FAS) pathway. mtFAS utilizes acetyl-CoA and Acyl Carrier Protein (ACP) to generate fatty acid chains at least 8 carbons in length. The fatty acids in mtFAS do not contribute to triglycerides or phospholipids, but instead generate octanoic acid for the synthesis of lipoic acid and long chain fatty acids (Wedan et al., 2024). Lipoic acid is a cofactor for Pyruvate Dehydrogenase (PDH) and α-Ketoglutarate Dehydrogenase (OGDH). Because these enzymes have key roles for carbon entry into the tricarboxylic acid (TCA) cycle, dysfunction of the mtFAS pathway can broadly impair mitochondria (Solmonson and DeBerardinis, 2018). Longer acyl chains generated from mtFAS have been implicated in electron transport chain (ETC) stability and mitochondrial translation (Dibley et al., 2020) and proposed as indispensable to maintain mitochondrial respiration (Nowinski et al., 2020; Tanvir Rahman et al., 2023).

Mitochondrial Trans-2-Enoyl Coenzyme A Reductase (MECR) catalyzes the last step in mtFAS to create acyl-ACP (Torkko et al., 2001). MECR has also been associated with iron and ceramide metabolism, and MECR-deficiency leads to an increase in both these pathways due to defective mitochondrial iron-sulfur cluster biogenesis (Dutta et al., 2023). Rare loss-of-function *MECR* mutations in humans were first described in 2016 with patients shown to have a mitochondrial deficiency disorder with dystonia and optic atrophy now termed Mitochondrial Enoyl CoA Reductase Protein-Assocoated Neurodegeneration (MEPAN) Syndrome (Heimer et al., 2016). However, the immune phenotypes in these patients have not yet been investigated in detail.

Here we employed unbiased approaches to identify essential lipid metabolic processes in CD4^+^ T cells. *In vivo* CRISPR/Cas9 screens performed using a custom lipid metabolism-based gRNA library identified mtFAS as having a previously undescribed role in T cells. We then generated *Mecr*^fl/fl^;*Cd4*^cre^ mice and found that while resting T cells were not affected, activated MECR-deficient T cells proliferated poorly and had reduced overall mitochondrial and oxidative metabolism. Intracellular iron was also elevated consistent with increased susceptibility to ferroptosis. MECR-deficiency decreased T cell fitness across multiple models including inflammatory bowel disease, airway inflammation, tumors, and infection. These findings show that although not essential for resting T cell viability, MECR promotes fitness of activated T cells.

## Results

### mtFAS genes regulate T cell fitness across multiple *in vivo* CRISPR/Cas9 screens

An *in vivo* CRISPR screening approach was employed in primary murine T cells to test the dependence of CD4^+^ and CD8^+^ T cells on lipid metabolism in models of inflammation (**Figure 1A**). A custom CRISPR/Cas9 library was constructed to target 47 lipid metabolism genes found in KEGG lipid metabolism database, with addition of 10 non-targeting negative control (NTC) and 2 positive control guides (*Tsc2* and *Rheb*) in a retroviral vector (**Supplementary Table 1**). CD4^+^ T cells were activated in Th17 promoting conditions, transduced with the lipid metabolism library, and adoptively transferred into *Rag1^-/-^* mice to induce Inflammatory Bowel Disease (IBD). Mice were sacrificed after disease developed and the T cells were isolated from the mesenteric lymph nodes (mLN) and the colonic lamina propria. The relative abundance of each guide RNA (gRNA) from T cells in the mLN and the lamina propria was determined by Next Gen Sequencing (NGS) and compared to gRNA frequencies prior to adoptive transfer. Interestingly, the sgRNAs for genes involved in mtFAS, including *Mecr*, *Oxsm*, and *Mcat,* were consistently depleted in T cells, suggesting a role for mtFAS in T cell fitness in the mLN and the lamina propria in the IBD model (**Figure 1B, C**).

**Figure 1:**
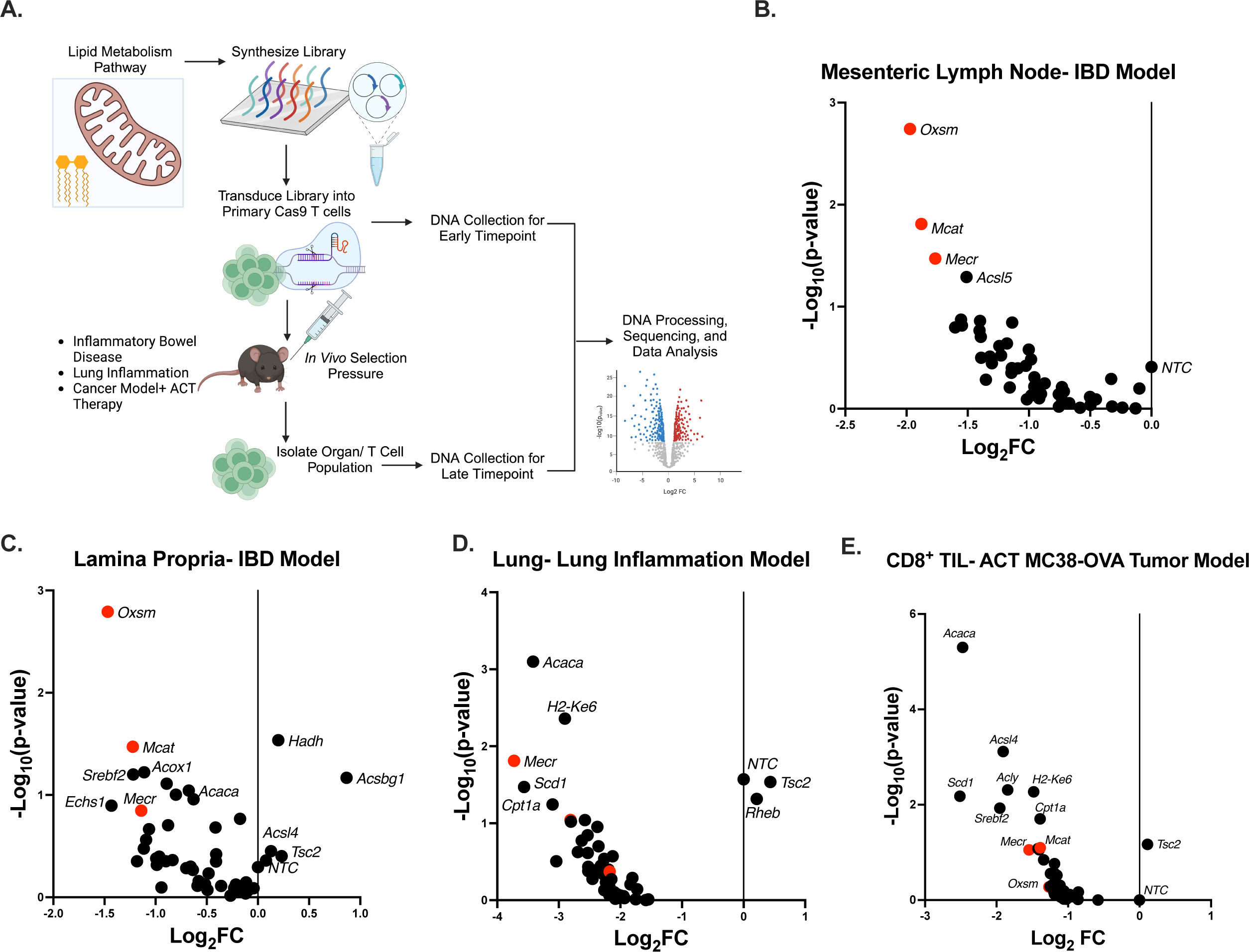
CRISPR/Cas9 *in vivo* screens identify mitochondrial fatty acid synthesis (mtFAS) genes. **a)** *In vivo* CRISPR/Cas9 screening protocol. **b-e)** Analysis of CRISPR/Cas9 Screens in an Inflammatory Bowel Disease model **(b,c**), lung inflammation **d)**, and CD8^+^ adoptive transfer therapy (ACT) MC38-OVA tumor model **(e)**. Statistical significance performed by MAGeCK. Panel (**d,e**) show the results of a representative experiment of 2 independent experiments. Panel b), n=6 biological replicates, Panel c), n=4 biological replicates, Panel d), n=3 biological replicates, Panel e), n= 4 biological replicates.

To validate these results, the same lipid metabolism CRISPR screen was performed in additional models. First, an antigen-specific model of lung inflammation was tested. Ovalubmin (OVA)-specific OT-II;Cas9 CD4^+^ T cells were activated and transduced with the lipid targeting gRNA library and adoptively transferred into *Rag1^-/-^* mice, which were then intranasally sensitized with OVA protein to promote lung inflammation. T cells recovered from the lung had depleted *Mecr*-targeting sgRNAs, showing the depletion is not disease specific **(Figure 1D)**. Second, a model of tumor adoptive transfer immunotherapy was utilized to test the dependence in CD8^+^ T cells on mtFAS. Transduced OT-I CD8^+^ T cells were transferred into MC38-OVA tumor-bearing *Rag1^-/-^* mice. After seven days, CD8 tumor-infiltrating T cells (TILs) were isolated from tumors for analysis. In this screen, *Mecr* and *Mcat* gRNAs were depleted in the antigen-specific CD8^+^ T cells in the tumor (**Figure 1E**). The depletion of sgRNAs of mtFAS in CRISPR screens across multiple *in vivo* models support a critical role for mtFAS and the genes *Mecr*, *Oxsm*, and *Mcat* in the fitness of both CD4^+^ and CD8^+^ T cells.

Because mtFAS has been poorly studied in immunity, *Mecr*, *Mcat*, and *Oxsm* expression was examined in immune cells. We first examined murine studies from available RNAseq datasets in T cell development and activation. While *Mcat* and *Oxsm* were expressed at low levels across thymic development, *Mecr* was dynamically regulated with high early expression and reduced expression in CD4^+^ CD8^+^ thymocytes (Yoshida et al., 2019) (**Supplemental Figure 1A**). *Mecr* expression, also, was increased after activation in mature murine antigen specific OT-I CD8^+^ T cells (**Supplemental Figure 1B**) (Madden et al., 2023). In LCMV Armstrong acute infection, *Mecr* expression initially increased after 6 days but then decreases (**Supplemental Figure 1C**). However, during a chronic LCMV clone 13 infection, chronically stimulated and exhausted T cells decreased *Mecr* expression over the course of 30 days (Doering et al., 2012). MECR protein expression was highly expressed in both CD8^+^ and CD4^+^ T cells, compared with MCAT and OXSM protein expression (Brenes et al., 2023) (**Supplemental Figure 1D**). Lastly, MECR protein levels increased after CD4^+^ and CD8^+^ T cell activation and this was dependent on Myc (**Supplemental Figure 1E**) (Marchingo et al., 2020).

### Acute *in vitro* disruption of *Mecr* with CRISPR/Cas9 does not alter T cell function

Naïve Cas9-transgenic CD4^+^ T cells were activated and skewed to Th1, Th17, or iTreg polarized subsets to test if loss of *Mecr* reduced T cell function in short-term *in vitro* conditions. T cells were transduced with sg*Mecr* or sgNTC and characterized by flow cytometry to evaluate the role of MECR on T cell differentiation and function (**Supplemental Figure 2A, B**). Neither IFNγ, IL-17a expression (**Supplemental Figure 2C, D**), nor Tbet (Th1), RORγt (Th17), or FoxP3 (iTreg) expression (**Supplemental Figure 2E-G**) were altered by acute MECR-deficiency. In addition, no differences were observed in general reactive oxygen species (DCFDA), total mitochondrial mass (Mitotracker), or mitochondrial membrane potential (TMRE) under these conditions (**Supplemental Figure 2H-J**). We hypothesized the maintenance of T cell function after *Mecr*-deletion may be due to with reliance on increased glycolysis *in vitro*. To test this, transduced cells were cultured in media containing glucose or galactose to enforce glycolysis or oxidative metabolism, respectively and stained for IFNγ to evaluate function of MECR-deficient T cells. While galactose media prevented aerobic glycolysis and decreased IFNγ, MECR-deficiency had no further effect on IFNγ levels (**Supplemental Figure 1K**). These data suggested the continued function of MECR-deficient T cells *in vitro* was not due to reliance glycolysis. Rather, MECR is either acutely non-essential or the timing of acute deletion and disruption of mitochondrial physiology was sufficiently delayed to mitigate *in vitro* phenotypes. Importantly, acute *in vitro* deletion was insufficient to account for the loss in fitness suggested by our *in vivo* screens.

### *Mecr*^fl/fl^; *Cd4*^cre^ mice have reduced CD4^+^ and CD8^+^ T cells *ex vivo*

Because chronic loss of MECR may be required to reveal the role of mtFAS in T cells, we developed a CD4^+^ and CD8^+^ T cell specific conditional *Mecr*-knockout mouse model (*Mecr*^fl/fl^; *Cd4*^cre^mice). Activated CD4^+^ T cells and MECR was confirmed to be reduced in *Mecr*-KO T cells by western blot **(Figure 2A)**. The percentage of thymocyte DN, DP, CD4 and CD8 SP cell populations were unchanged consistent with late thymic Mecr deletion (**Supplemental Figure 3A-D**). Total numbers of live splenocytes between control cre-(WT) and cre+ (*Mecr*-KO) T cells (**Supplemental Figure 3E**) as well as total number of peripheral CD4^+^ and CD8^+^ T cells were unchanged (**Supplemental Figure 3F, G**). However, the percentages of CD4^+^ and CD8^+^ T cells in the spleen were significantly reduced (**Figure 2B, C**). Interestingly, the CD4^+^ T cell compartment of MECR-deficient mice skewed toward and increased proportion of naïve cells and reduced fraction of effector (CD62L^-^ CD44^+^) and central memory (CD62L^+^ CD44^+^) CD4^+^ T cells (**Figure 2D-G**). MECR-deficient T cells also had decreased percentages of Tbet^+^ cells and IFNγ^+^ CD4^+^ T cells (**Figure 2H, I**), suggesting a defect in Th1 effector T cells. However, Treg populations appeared normal (**Supplemental Figure 3H, I**). CD4^+^ and CD8^+^ T cell percentages were also decreased in the liver (**Figure 2J, K**), showing that *Mecr*-KO T cells had a fitness disadvantage in tertiary immune landscapes. MECR, therefore, was not critical for naïve T cells or their homing in lymph nodes and the spleen, but was required for full activation of effector-memory populations and tissue homing.

**Figure 2:**
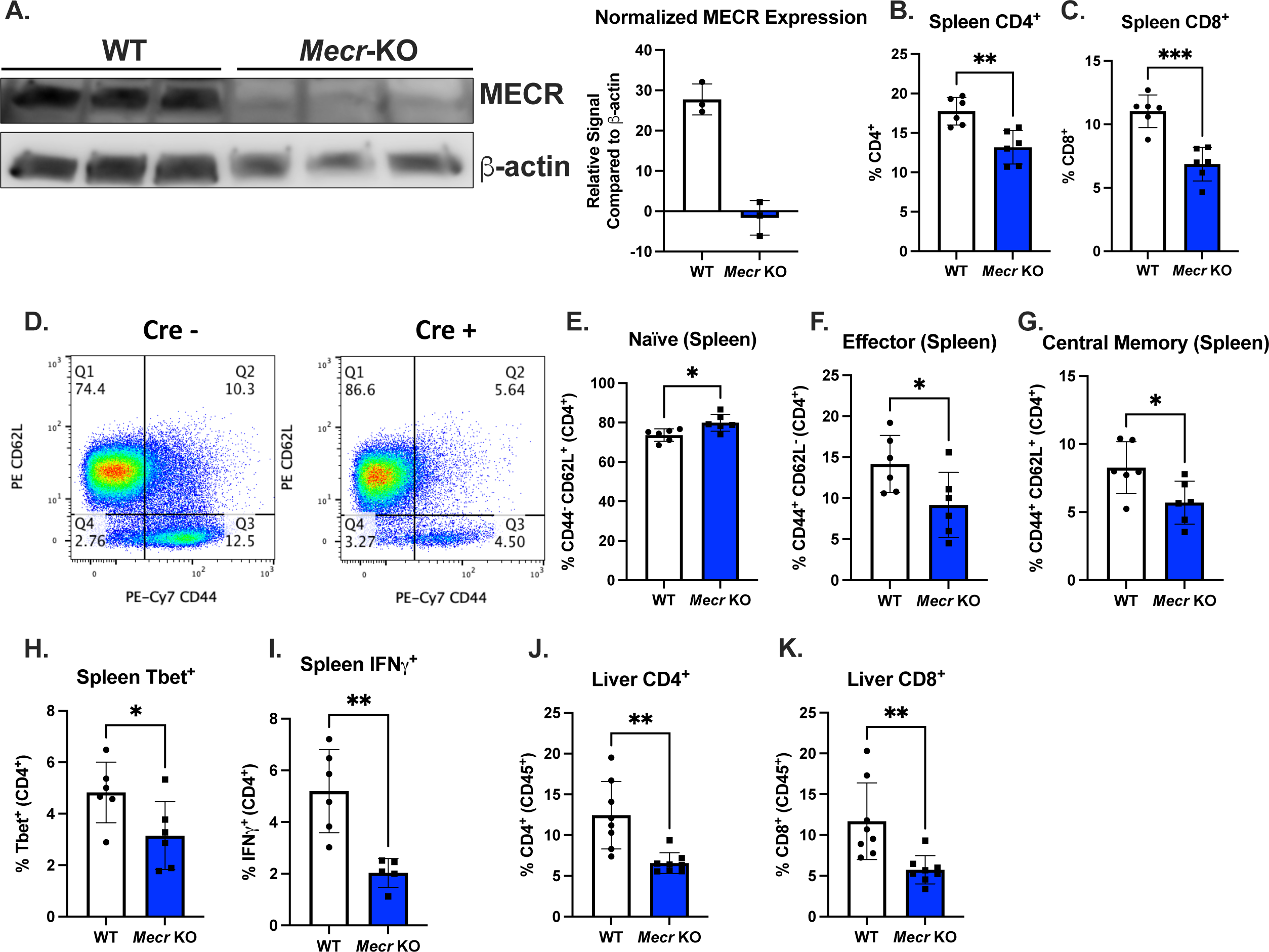
*Mecr*^fl/fl^; *Cd4*^cre^ mice have reduced CD4 and CD8 T cells *ex vivo*. **a)** Western blot of MECR expression in activated CD4^+^ T cells. **b)** Percentage of CD4^+^ T cells in the spleen measured by flow cytometry**. c)** Percentage of CD8^+^ T cells in the spleen. **d-g)** Naïve, effector, and central memory CD4^+^ T cells in the spleen. **h, i)** Th1 function percentage of IFNγ^+^ and Tbet. **j, k)** Percentage of CD4^+^ and CD8^+^ in the liver. Panels (b-k show results from two pooled independent experiments. Each data point represents a biological replicate and error bars show standard deviation. All Statistical significance performed by unpaired t tests. (* p<0.05, ** p<0.01, *** p<0.001, **** p<0.0001).

### MECR is required for efficient CD4^+^ T cells proliferation and differentiation

We next tested the effects of chronic loss of MECR on CD4^+^ T cell proliferation, survival, and differentiation. CD4^+^ T cells were isolated from *Mecr*^fl/fl^; *Cd4*^cre^ mice and activated with anti-CD3, anti-CD28, and IL2. After three days, *Mecr*-KO T cells had significantly reduced proliferation as measured by the cell division index of cell trace violet (CTV) dilution compared to the cells from WT mice (**Figure 3A).** The mTORC1 pathway responds to available nutrients and has multiple downstream metabolic mechanisms to promote cell growth and proliferation by increasing nucleotide, protein, and lipid synthesis. To test if the effects of MECR on proliferation are linked to mTORC1 activity, we stained activated *Mecr*^fl/fl^; *Cd4*^cre^ CD4^+^ T cells cells for phospho-S6, S235/236, as a measurement of mTORC1 activity. *Mecr*-KO cells had significantly reduced P-S6 MFI compared to controls (**Figure 3B**). Therefore, MECR is essential for mTORC1 activity.

**Figure 3:**
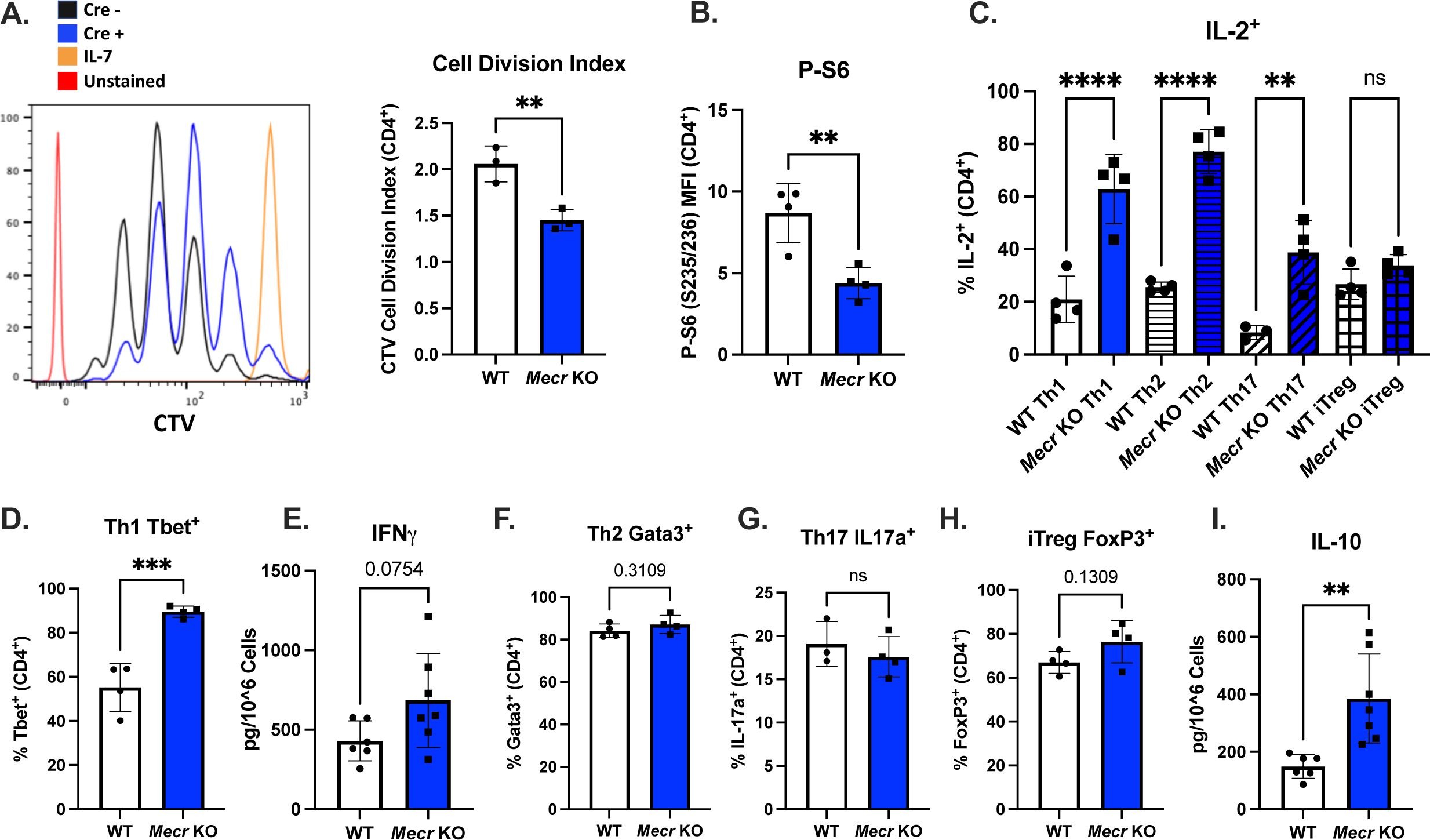
MECR is required for T cell proliferation and differentiation. **a)** Cell division index of activated CD4^+^ T cells stained with cell trace violet (CTV) measured by flow cytometry. **b)** phospho-S6 MFI of activated CD4^+^ T cells. **c)** Percentage of IL-2^+^ T cells in each T cell subset. **d-h)** Differentiated naïve CD4^+^ T cell transcription factors of Th1 Tbet **(d)**, Th2 Gata3 **(f)**, Th17 RORγt **(g)**, and iTreg FoxP3 **(h)** and cytokine production of Th1 IFNγ **(e)**, and iTreg IL-10 **(i)**. Panels (a) show representative results from three independent experiments. Panels (b, c, d, f, g, h) show representative results from two independent experiments and panels (e, i) show pooled results from two independent experiments. Each data point represents a biological replicate and error bars show standard deviation. All Statistical significance performed by unpaired t tests. (* p<0.05, ** p<0.01, *** p<0.001, **** p<0.0001).

To test if MECR impacts T cell differentiation *in vitro*, naïve CD4^+^ T cells were isolated from *Mecr*^fl/fl^; *Cd4*^cre^ mice and activated with skewing cytokines for Th1, Th2, Th17, or iTreg polarization. After three days, the ability for cells to differentiate was measured by lineage specific transcription factors and cytokines along with CTV. Proliferation was reduced in all MECR-deficient subsets, with Th17 and iTregs the most impacted followed by Th1 then Th2 (**Supplemental Figure 3J**). Interestingly, the percentage of IL-2^+^ cells was increased by MECR-deficiency in the proinflammatory Th1, Th2, and Th17 subsets, suggesting the loss of Mecr may impact the differentiation of one or more of these cell subsets (**Figure 3C**). To test this, we quantified lineage specific transcription factors and cytokines for CD4 T cell subsets along with CTV. Tbet^+^ T cells were significantly increased and there was a trending increase in IFNγ production as measured by ELISA in MECR-deficient T cells, which indicate skewing towards a Th1 phenotype (F**igure 3D, E**). In contrast, both the Th2 associated transcription factor Gata3 and the Th17 associated cytokine IL-17a were not significantly different in *Mecr*-KO cells, indicating acute loss of MECR does not affect the differentiation of CD4^+^ T cells into the Th2 or Th17 subset (**Figure 3F, G**). In addition, the iTreg associated transcription factor FoxP3 showed a modest elevated trend with MECR-deficiency (**Figure 3H**) and IL-10 production was increased in MECR-deficient iTregs as measured by ELISA (**Figure 3I**). These data show that Mecr is required for effective CD4^+^ proliferation and survival and affects CD4^+^ subsets differentially, with a possible role in differentiation of Th1 and iTreg cells.

### MECR-deficiency reduces mitochondrial function and oxidative metabolism

*Mecr*-KO skeletal myoblast cells have disrupted mitochondrial function (Nowinski et al., 2020). To test if this phenotype was similar in T cells, activated *Mecr*^fl/fl^; *Cd4*^cre^ CD4^+^ T cells were stained for mitochondrial function, including mitochondrial mass (mitotracker), mitochondrial membrane potential (TMRE), and mitochondrial reactive oxygen species-mtROS (Mitosox). Mitotracker and TMRE were significantly increased in MECR-deficient cells (**Figure 4A, B**) suggesting increased mitochondrial size and dysregulation of the ETC proton gradient. In addition, mtROS was significantly increased in activated *Mecr*-KO CD4^+^ T cells (**Figure 4C**). However, total ROS measured by DCFDA was unchanged, suggesting a mitochondrial specific effect or compensatory cytosolic adaptations (**Figure 4D**).

**Figure 4:**
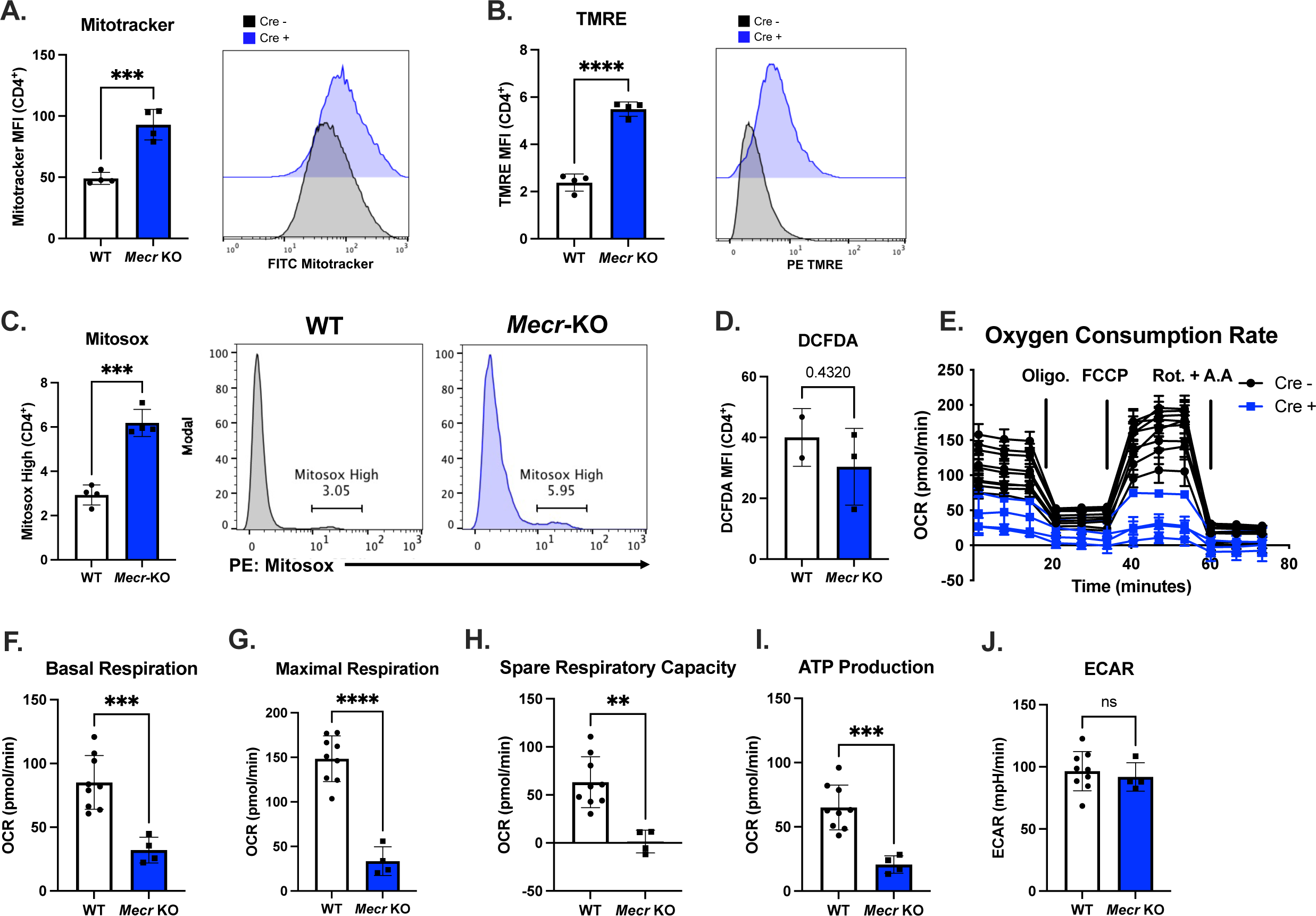
*Mecr*-KO causes reduced mitochondrial function and oxidative metabolism. **a-d)** Mitochondrial stains in activated CD4^+^ T cells from WT and *Mecr*-KO mice, Mitotracker **(a)**, TMRE **(b)**, Mitosox **(c)**, DCFDA **(d)**. **e)** Extracellular flux analysis on activated CD4^+^ T cells. **f, g)** Basal and maximal respiration. **h)** Spare respiratory capacity. **i)** ATP production. **j)** Extracellular acidification rate (ECAR). Panels (a-d) show representative results from two independent experiments and each data point represents a biological replicate. Panels (e-j) show results from two pooled independent experiments. Each data point represents a biological replicate with averaged n>3 technical replicates. Error bars show standard deviation. All Statistical significance performed by unpaired t tests. (* p<0.05, ** p<0.01, *** p<0.001, **** p<0.0001).

Reduced ETC stability with MECR-deficiency may reduce oxidative metabolism caused by reduced ETC stability, as acyl-ACP-dependent LYRM proteins that maintain ETC complexes can decrease in the absence of mtFAS (Nowinski et al., 2020). To test if similar respiration defects were observed in *Mecr*-KO CD4^+^ T cells, we conducted a mitochondrial stress test to analyze T cell mitochondrial function (**Figure 4E)**. Indeed, the basal and maximal respiration of the MECR-deficient CD4^+^ T cells were significantly reduced, spare respiratory capacity was nearly eliminated, and ATP production was reduced when compared to wildtype counterparts (**Figure 4F-I**). In contrast, the basal extracellular acidification rate (ECAR), which reflects the activity of aerobic glycolysis was unchanged in MECR-deficient T cells (**Figure 4J**). With dysregulated mitochondria function and respiration, these data show that MECR is needed for efficient ATP-linked mitochondrial respiration.

### CD4^+^ T cells lacking MECR have impaired TCA cycling and ETC metabolism

To further evaluate mitochondrial function in MECR-deficient CD4^+^ T cells, ETC complex proteins were analyzed by immunoblot. ETC complexes I and II were most reduced in *Mecr*-KO T cells (**Figure 5A**), confirming ETC defects. The mtFAS product acyl-ACP can also be used to synthesize lipoic acid, which is essential for the activity of Pyruvate Dehydrogenase (PDH) and α-Ketoglutarate Dehydrogenase (OGDH). There was an almost complete reduction in lipoic acid on PDH complex DLAT and OGDH complex DLST in MECR-deficient T cells (**Figure 5A**). MECR, therefore, is needed for ETC complex assembly in T cells.

**Figure 5:**
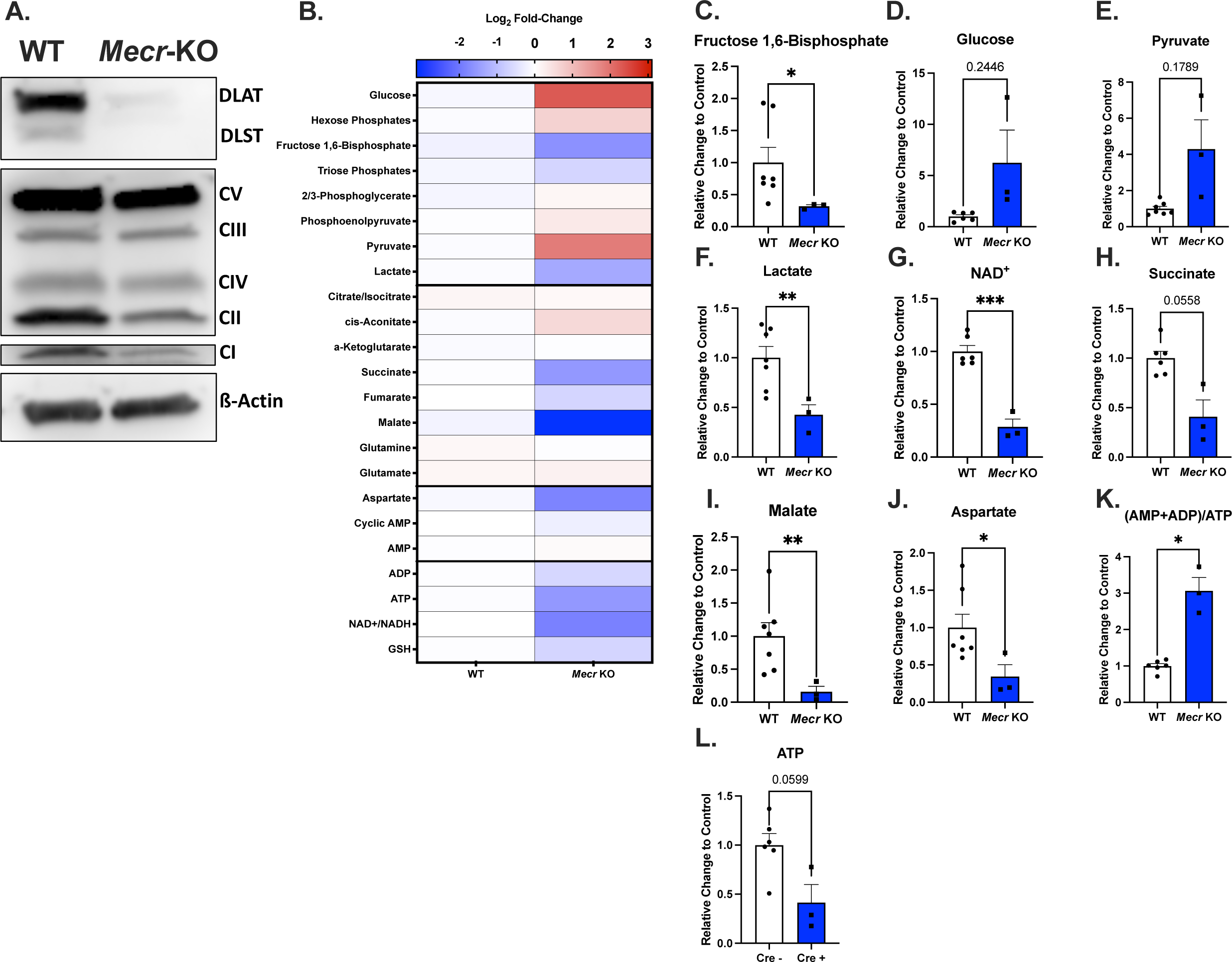
*Mecr*-KO T cells have reduced oxidative metabolism. **a)** Western blot of activated CD4^+^ T cells blotted for lipoic acid on DLAT and DLST, OXPHOS complexes I-V, and ß-actin. **b)** Heatmap of metabolites from metabolomics. **c-k)** Quantified relative change of metabolites in activated CD4^+^ T cells with *Mecr*-KO compared to WT cells, fructose 1,6-bisphosphate **(c)**, glucose **(d)**, pyruvate **(e)**, lactate **(f)**, NAD^+^ **(g)**, Succinate **(h)**, malate **(i),** aspartate **(j)**, (AMP+ADP)/ATP **(k)**, ATP **(l)**. Panels (a-l) show results from one independent experiment. Each data point represents a biological replicate and error bars show standard error of the mean. Statistical significance panels (c-k) performed by Welch’s t test. (* p<0.05, ** p<0.01, *** p<0.001, **** p<0.0001).

We next analyzed steady-state abundance of TCA intermediates in activated CD4^+^ T cells from *Mecr*^fl/fl^; *Cd4*^cre^ mice to better understand the role of MECR in T cell mitochondrial metabolism (**Figure 5B**). While fructose 1,6-bisphosphate was significantly reduced (**Figure 5C**), intracellular glucose and pyruvate showed trends towards increased abundance (**Figure 5D, E)**. Conversely, lactate and NAD^+^ were decreased (**Figure 5F, G**) and succinate trended downward (**Figure 5H**) suggesting impaired conversion of α-ketoglutarate to downstream succinate along with impaired conversion of NAD^+^ for use by the ETC (**Figure 5H**). Malate was also significantly decreased in MECR-deficient cells (**Figure 5I**), potentially due to the shuttling of malate into the ETC through the malate-aspartate shuttle before conversion into oxaloacetate. In that same vein, aspartate was also significantly decreased, concordant with a requirement of oxaloacetate as an electron acceptor in the ETC complex through the malate-aspartate shuttle (**Figure 5J**). Lastly, the ATP energy ratio ((AMP+ADP)/ATP) in the Mecr-deficient CD4 T cells was elevated compared to controls and ATP was reduced (**Figure 5K, L**), suggesting MECR-deficiency in CD4^+^ T cells reduces the ability of ETC1 to produce protons that fuel ATP synthesis. MECR, therefore, maintains the oxidative and ETC metabolism of activated CD4^+^ T cells.

### *Mecr*-knockout T cells accumulate iron and are susceptible to ferroptosis

MECR-deficient cells from *Drosophila* and human MEPAN patient fibroblasts accumulate excess iron and ceramides due to defective Fe-S cluster biogenesis in the mitochondria (Dutta et al., 2023). T cell subsets have differential requirements for iron, with Th1 being more dependent on iron than Th2 (Thorson et al., 1991), as well as Th17 cells requiring iron for differentiation and IL-17a production (Li et al., 2021). Further, blocking the transferrin receptor (CD71) for iron uptake alters mitochondrial function and OXPHOS in murine and human T cells (Voss et al., 2023). We hypothesized therefore that *Mecr*-KO T cells may have dysregulated iron metabolism that impairs mitochondrial respiration. CD4^+^ T cells from *Mecr*^fl/fl^; *Cd4*^cre^ mice were activated for three days and cellular labile ferrous iron (II), mitochondrial iron, and CD71 expression were measured. All were significantly increased in MECR-deficient T cells compared to control cells (**Figure 6A-C**). In addition, lipid peroxidation, as measured by C11 BODIPY was also increased in T cells lacking MECR (**Figure 6D)** and there was a significant increase in the percent of cell death in activated T cells (**Figure 6E, F**).

**Figure 6:**
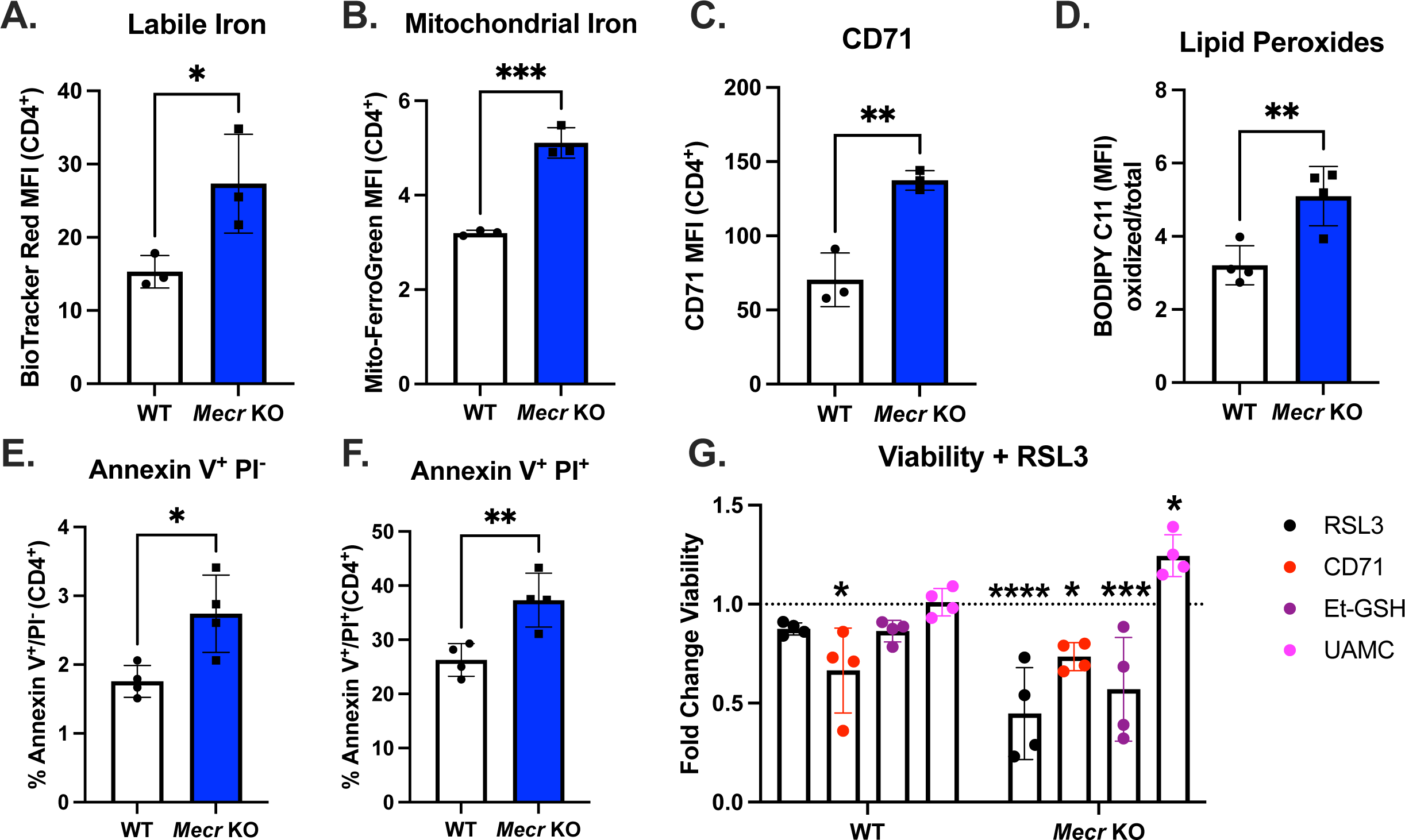
*Mecr*-KO T cells accumulate iron which causes increased sensitivity to ferroptosis. **a-c)** Activated CD4^+^ T cells stained for labile (2+) iron **(a)**, mitochondrial iron **(b)**, CD71 **(c),** and C11 BODIPY **(d)**. **e, f)** Percentage of apoptosis **(e)** and cell death by apoptosis **(f)** of activated T cells. **g)** Viability of activated T cells with treatment of RSL3. Panels (a-c) show representative results from three independent experiments. Panel d, g shows results from one independent experiment. Panels (e,f) show results from two independent experiment. Each data point represents a biological replicate and error bars show standard deviation. Panels (a-f) statistical significance performed by unpaired t tests. Panel (g) statistical significance performed by two-way ANOVA with uncorrected Fisher’s LSD test. (* p<0.05, ** p<0.01, *** p<0.001, **** p<0.0001).

Because of the increased iron and cell death, we hypothesized that the observed increase in cell death may be caused by ferroptosis. To test this, activated WT and *Mecr*-KO T cells were treated with mouse anti-CD71 blocking antibody or isotype control for three days. As shown previously, anti-CD71 treatment during activation lowers iron uptake by internalizing CD71 receptors in murine T cells and reduces viability (Voss et al., 2023). After 3 days of activation, all cell treatments were challenged with the GPX4 inhibitor RSL3 to induce ferroptosis and cell death. Cells were also pre-treated with cell-permeable glutathione (Et-GSH) or UAMC-3203 (Devisscher et al., 2018) to inhibit ferroptosis. *Mecr*-KO cells had reduced viability when treated with ferroptosis inducer RSL3 compared to WT cells, demonstrating increased susceptibility to ferroptosis (**Figure 6G**). Importantly, *Mecr*-KO cell viability was rescued when treated with the specific ferroptosis inhibitor UAMC-3203, but not anti-CD71 or Et-GSH which nonspecifically lower iron content or increase antioxidant capacity, respectively. These data show that *Mecr*-KO cells accumulate iron and lipid peroxidation, which may contribute to increased cell death by ferroptosis.

### Loss of MECR reduces T cell survival and function in multiple *in vivo* models

Because *Mecr* was consistently depleted across several *in vivo* CRISPR screens and promoted mitochondrial metabolism of activated T cells, we hypothesized that *Mecr*-KO T cells have reduced fitness and reduced transcription factors and cytokines when directly tested in *in vivo* disease models. Th17 skewed Cas9-transgenic CD4^+^ T cells were transduced with non-targeting control (NTC) sgRNA in a GFP retroviral vector or a *Mecr*-targeting sgRNA in a BFP vector shown to induce knockout (**Supplemental Figure 4A**). Cells were mixed at a 1:1 ratio (**Figure 7A**) and injected into *Rag1*^-/-^ mice to induce IBD. After 8 weeks the frequency of *Mecr*-KO CD4^+^ T cells was significantly reduced in the spleen, mesenteric lymph nodes (mLN), lamina propria, intra-epithelial lymphocytes, and the skin compared to control cells (**Figure 7B-G**), confirming that MECR loss reduces survival or recruitment of activated T cells. *Mecr*-KO cells had increased surface expression of CD62L and decreased CD44 (**Figure 7H, I**), suggesting the remaining T cells have reduced effector function.

**Figure 7:**
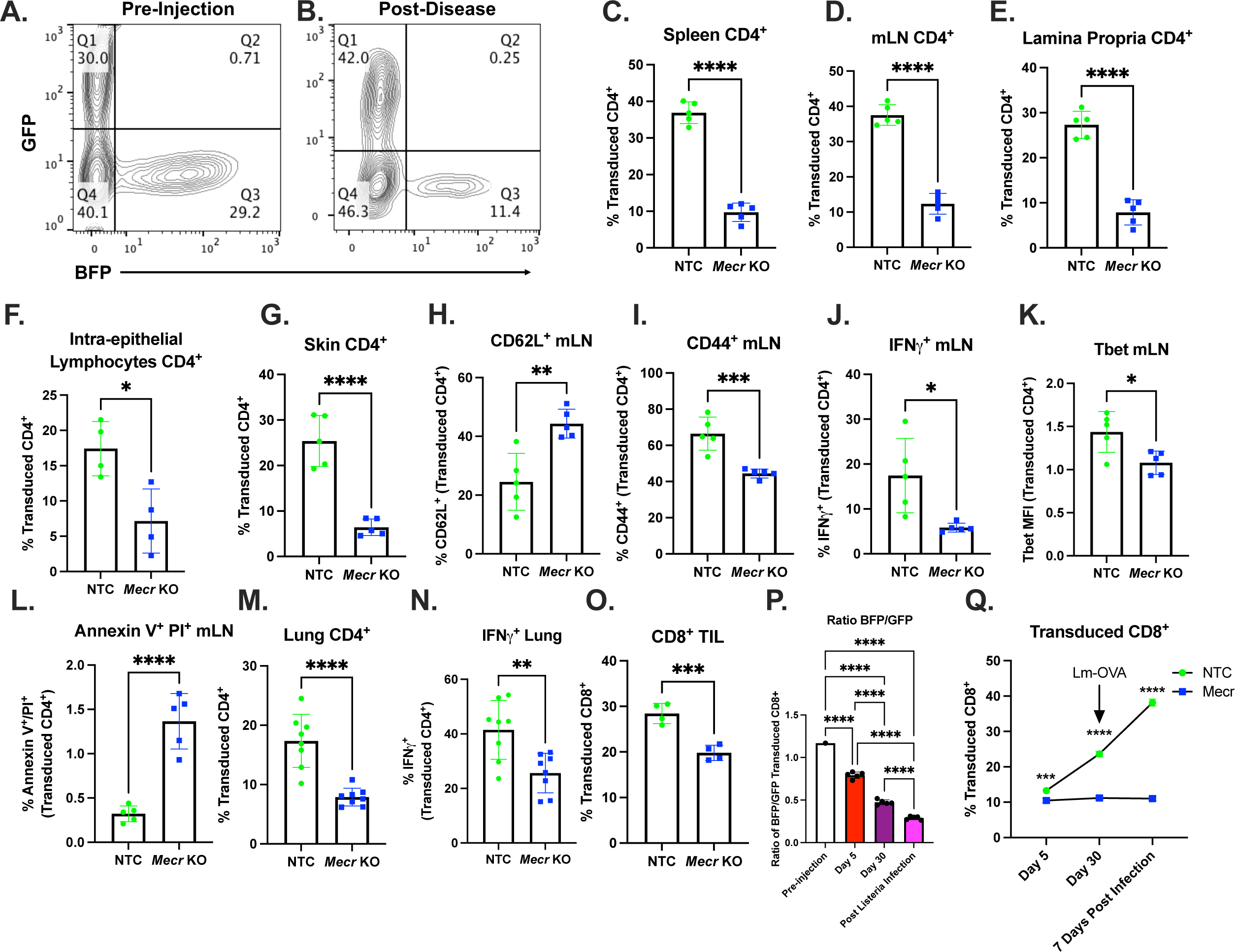
*Mecr*-KO causes reduced survival in multiple *in vivo* models. **a)** 1:1 mixed sgNTC T cells (GFP) and sg*Mecr* (BFP) pre-injection. **b)** GFP^+^ and BFP^+^ T cells of CD4^+^ T cells post-IBD development. **c-g)** Percentage of transduced CD4^+^ expressing GFP (NTC) or BFP (*Mecr*-KO) in the spleen **(c)**, mesenteric lymph node (mLN) **(d)**, lamina propria **(e)**, intra-epithelial lymphocytes **(f)**, and skin **(g)**. **h)** CD62L^+^ of transduced CD4^+^ T cells. **i)** CD44^+^ of transduced CD4^+^ T cells. **j)** IFNγ^+^ of transduced CD4^+^ T cells. **k)** Tbet mean fluorescence intensity (MFI) of transduced CD4^+^ T cells. **l)** Annexin V^+^ Propidium Iodide (PI)^+^ of transduced CD4^+^ T cells. **m, n)** lung inflammation model percentage of transduced CD4^+^ T cells **(m)**, and IFNγ^+^ of transduced CD4^+^ T cells **(n)**. **o)** Percentage of transduced CD8^+^ T cells in the MC38-OVA tumor. **p, q)** Listeria-OVA antigen-specific CD8 model ration of BFP:GFP of transduced CD8^+^ T cells from the spleen **(p)**, and percentage of transduced CD8 T cells in the spleen over time **(q)**. Panels (a-l) show representative results from >5 independent experiments. Panels (m,n) show representative results from one of two 2 independent experiments. Panel (o) shows results from one independent experiment. Panels (p, q) representative of 2 independent experiments. Each data point represents a biological replicate and error bars show standard deviation. All Statistical significance performed by unpaired t tests with the exception of (p) with a one-way ANOVA with a Dunnett’s multiple comparison. (* p<0.05, ** p<0.01, *** p<0.001, **** p<0.0001).

To further investigate *Mecr*-KO T cell function and phenotypes, cytokine and transcription factors were quantified. Th17-skewed cells adopted a Th1 phenotype in colitis, as previously published (Harbour et al., 2015). With this IBD model, *Mecr*-KO T cells had a significantly reduced percentage of IFNγ^+^ T cells compared to the NTC T cells as well as reduced Tbet expression **(Figure 7J, K)**, suggesting impaired Th1 function. Interestingly, there was no difference in RORγt expression, a marker of Th17 phenotypes (**Supplemental Figure 4B**), although there was a trend for decreased percentages of IL-17a^+^ IFNγ^+^ and IL-17a^+^ IFNγ^-^ T cells (**Supplemental Figure 4C-E**). In addition, *Mecr*-KO T cells exhibit increased cell death by apoptosis (Annexin V^+^ PI^+^) compared to NTC transduced T cells (**Figure 7L**). The antigen-specific CD4^+^ OT-II lung inflammation was also used to confirm a loss of fitness in *Mecr*-KO cells using the T cell competition model. After disease development and lungs were harvested, *Mecr*-KO cells were also significantly reduced in the lung-inflamed tissue in addition to the *Mecr*-KO T cells expressing reduced percentages of IFNγ^+^, showing reduced Th1 CD4^+^ T cell function (**Figure 7M, N**).

The effects of MECR-deficiency on CD8^+^ T cells *in vivo* were next tested using the competitive transfer approach with CRISPR/Cas9 single-guide knockouts. We first used an adoptive cell transfer therapy cancer model in which Ovalbumin (OVA)-specific OT-I CD8^+^ T cells are transferred into recipients bearing MC38-OVA tumors. *Mecr*-KO cells accumulated less well in MC38-OVA tumors, although it was unclear if this was due to impaired recruitment, proliferation, or survival (**Figure 7O**). Finally, a CD8^+^ listeria-OVA infection 1:1 competition model was used to evaluate the loss of fitness in *Mecr*-KO CD8^+^ T cells. Comparing pre-adoptive transfer, days 5, day 30, and post-listeria-OVA infection, the ratio of *Mecr*-KO to NTC T cells was significantly reduced at initial expansion on day 5 and in the memory population on day 30 post-infection (**Figure 7P**). One week after rechallenge *Mecr*-KO T cells remained lower in abundance on day 37 (**Figure 7Q**). In additon, the total numbers of live transduced NTC CD8^+^ T cells expanded in this model, while the *Mecr*-KO cells did not (**Supplemental Figure 4F**). These data show that the loss of MECR causes reduced CD4^+^ and CD8^+^ T cell survival and function *in vivo*.

## Discussion

T cells reprogram lipid synthesis upon activation to support growth and signaling. In this study, we applied an unbiased CRISPR/Cas9 *in vivo* screening approach to identify key lipid metabolizing enzymes in T cells *in vivo* and found mtFAS and *Mecr* to be critical for T cell recruitment and persistence in inflammation. Our results show that MECR regulates CD4^+^ T cell function and oxidative metabolism. Given the role of MECR in mtFAS, it was surprising that naïve *Mecr*-KO T cells were largely unaffected. Nevertheless, *Mecr*-KO cells exhibited significant fitness disadvantages in models of IBD and lung inflammation. but did retain some functionality. In contrast, it is interesting to consider the lethal effects of mutating this pathway, as global knockout of *Mecr* is embryonically lethal in mice and overexpression causes cardiac dysfunction (Chen et al., 2009; Nair et al., 2017).

MECR had context-specific effects on CD4^+^ T cell subsets, with Th1 cells increased in acute *in vitro* assays but decreased IFNγ observed in chronic *in vivo* IBD models. In contrast, natural Tregs were unaffected, however, iTreg differentiation showed increased IL-10 production and trending increase in FoxP3 expression in *Mecr*-KO cells. This finding was somewhat unexpected, as we hypothesized that MECR would negatively affect Tregs due to their reliance on fatty acid oxidation and OXPHOS for energy, as compared to aerobic glycolysis and fatty acid synthesis in effector T cells (Michalek et al., 2011). This may be explained in part by impaired mTORC1 activity as measured by phospho-S6. It has been previously shown that the differentiation of Th1 and Th17 cells are regulated by mTORC1 while Th2 are regulated by mTORC2 (Delgoffe et al., 2011), and T cells treated with an mTORC1 inhibitor, rapamycin, have reduced effector T cell proliferation and function (Zhang et al., 2023).

In contrast to the results *in vivo* and *ex-vivo* utilizing *Mecr*^fl/fl^; *Cd4*^cre^ mice which showed a clear role for MECR, acute deletion of *Mecr* resulted in no significant differences *in vitro*. This may be due to prolonged persistence of MECR protein after CRISPR deletion, stability of the ETC complexes, long half-life of ETC complexes, or metabolic differences *in vitro* versus *in vivo.* It was possible that T cells obtained lipoic acid from serum in cell culture media, but previous studies found that supplementing with lipoic acid does not rescue ETC assembly and lipoic acid must be made endogenously to post-translationally modify enzymes as they are synthesized and fold into protein complexes (Nowinski et al., 2020; Solmonson and DeBerardinis, 2018). Overall, our data are most consistent with a long half-life and persistence of lipoic acid-modified proteins and ability of some cells to persist with reduced ETC activity.

MECR-deficient skeletal myoblasts were previously shown to have impaired basal respiration and low spare respiratory capacity. This was in part due to reduced acyl-ACP in complex with LYRM proteins to help form ETC complexes. Consistent with this, ETCs I, II, and IV were almost completely absent in MECR-deficient skeletal myoblasts (Nowinski et al., 2020). Impaired oxidative metabolism and ETC deficiency were also observed in MECR-deficient T cells. Reduced mitochondrial respiration, as seen by the mitochondrial stress test, agrees with other studies (Nowinski et al., 2020; Webb et al., 2023) in addition to reduced lipoic acid and ETC complexes I, II, and IV that LYRM proteins help stabilize. However, depending on mutations in the pathway, lipoic acid synthesis may be retained, as shown by a case study with a patient harboring a mutation in *MCAT* (Webb et al., 2023), confirming that lipoic acid is not the only biologically important product generated by the mtFAS pathway and supplementing lipoic acid does not rescue mitochondrial ETC assembly defects. We did not observe increased basal extracellular acidification rate. However, the metabolomics shows trending increases in glucose and pyruvate with a significant reduction in fructose 1,6-bisphosphate. With our metabolomics data, we saw reduced TCA cycle intermediates and defective ETC complexes. Increased pyruvate may be due to lipoic acid being a cofactor for PDH and may cause an accumulation due to impaired acetyl-CoA biosynthesis. TCA cycle intermediates succinate and malate were reduced in *Mecr*-KO cells and there was an accumulation of NAD^+^ and reduced production of ATP. Reduced aspartate also suggests a defect in the function of the ETC. Cumulatively, depletion of ETC interacting TCA intermediates in Mecr-KO CD4^+^ T cells support the data on ETC dysregulation. Our data also suggest that excess mtROS in MECR-deficient T cells may cause reduced T cell function and that the reduced mitochondrial function could contribute to reduced T cell memory and dysfunction *in vivo*.

MECR-deficient T cells also have increased levels of labile iron, mitochondrial iron, and CD71. This correlates well with *Mecr*-KO in *Drosophila* and fibroblast MEPAN patient samples also having an accumulation of intracellular iron (Dutta et al., 2023). Consistent with elevated iron accumulation, *Mecr*-KO CD4^+^ T cells were more susceptible to ferroptosis, which may contribute the fitness disadvantages observed *in vitro* and *in vivo*. However, because mtFAS has many products contributing to different pathways, we anticipated we could not fully rescue CD4^+^ T cell function by solely targeting ferroptosis. Regardless, these findings corroborate our oxidative metabolism data, in which ETC complexes I and II, which require those clusters, are reduced. As iron is essential in T cells and can control mitochondrial function and T cell activation through Fe-S clusters and mtROS (Sena et al., 2013), this is another potential way in which Mecr-KO is causing reduced activation and function of T cells.

Interestingly, mitochondrial disease patients often have immune dysfunction, with recurrent upper respiratory infections that can lead to sepsis, pneumonia, and other health complications such as gastroenteritis and leukopenia (Kapnick et al., 2018). While we would expect MEPAN patients to exhibit similar immune dysfunction base on our data, no previous case report has evaluated immune function, possibly due to the rarity of the patients and overall complexity of the disease presentation. Our data show that MECR and the mtFAS pathway are vital for the function of effector T cells. This study may provide opportunities to further investigate mtFAS and its immunological impact in MEPAN patients and suggest that MECR and mtFAS may be a potential target to treat inflammatory diseases.

## Materials and Methods

### Cell Lines

PlatE cells (Cell BioLabs RV-101) were used for transfection and retroviral production of sgRNA. PlatE cells for passaging were cultured with 1ug/mL puromycin (Gibco A11138-03) and 10ug/mL blasticidin (Gibco A11139-03). MC38-OVA tumor cells were generated as previously described (Madden et al., 2023). Cell lines were mycobacterium tested and fingerprinted.

### Mice

*Rag1^-/-^* (Jackson Labs #002216), Cas9 (Jackson Labs 024858), *Cd4*^cre^ (Jackson Labs 022071), OT-I (Jackson Labs #003831), and OT-II (Jackson Labs #004194) mice were obtained from Jackson Laboratory. *Rag1^-/-^* mice were used for *in vivo* CRISPR/Cas9 adoptive transfer recipients and Cas9 mice were used for CRISPR/Cas9 experiments to isolate T cells that express Cas9. *Mecr*^fl^ mice were obtained from Genpharmatech (T007874) and crossed to *Cd4*^cre^ mice to generate *Mecr*^fl/fl^; *Cd4*^cre^. Genotypes were validated with DNA sequencing through Transnetyx. For IBD models, 8-20 week old male and female *Rag1^-/-^* mice were used and euthanized when humane endpoint was reached (>20% weight loss). For injectable tumor models, 8-20 week old male and female *Rag1^-/-^* mice were used. Mice were euthanized when humane endpoint was reached (2cm dimension, ulceration, weight loss >10%). Both male and female mice were utilized in experiments. All mouse procedures were performed under Institutional Animal Care and Use Committee (IACUC)-approved protocols from Vanderbilt University Medical Center and conformed to all relevant regulatory standards.

### T Cell Isolation

Mouse spleens and lymph nodes were dissociated with a syringe, passed through a 70μm filter, and lysed using ACK lysis buffer. CD4^+^ T cells were isolated using the Stem Cell Easy Sep Mouse CD4^+^ T cell Isolation Kit (Stem Cell 19852) according to the manufacturer’s protocol. For CD4^+^ naïve cell isolation, the Miltenyi Naïve CD4^+^ T Cell Isolation Kit (Miltenyi 130-104-453) was used according to the manufacturer’s protocol. Isolated Cells were resuspended at 1 million cells/mL in complete RPMI 1640 (Corning, MT10040CV) media supplemented with 10% FBS (Avantor, 97068-085),100U/mL penicillin/streptomycin (Gibco, 15140122), 10mM HEPES (Thermo Fisher, 15630080), 2mM glutamine (Media Tek, 25030081), and 50µM 2-mercaptoethanol (Gibco, 21985023) and cultured at 37°C with 5% CO_2_. For galactose assay, galactose (Sigma Aldrich, G5388-100G) was used at equimolar to glucose.

### Isolation of cells from the liver and skin

Euthanized mice were perfused through the right ventriculus with 30 mL of ice-cold PBS. The liver and ears were dissected, washed in PBS, minced to small pieces with scissors and digested for 30 min at 37°C in DMEM + 0.5% BSA + 0.1% collagenase D. Cell suspension was filtered through a 70 μm strainer and cell debris was removed by centrifugation at 400 g for 20 min on 30% Percoll gradient loaded on top of 80% Percoll solution in PBS.

### CRISPR Screen and sgRNA Generation

The targeted lipid metabolism library was generated by referencing the KEGG Pathway Database (https://www.genome.jp/kegg/pathway.html) in addition to genes found in literature searches (**Supplemental Table 1**). Four sgRNAs per each gene found in the CRISPR Knockout Pooled Library (Brie) along with 10 non-targeting controls (NTC) for a total of 47 genes and 198 targets. The gRNA sequences flanked by the following adaptor sequences were purchased as an oligo pool from Twist Bioscience: GGAAAGGACGAAACACCGXXXXXXXXXXXXXXXXXXXXGTTTTAGAGCTAGAAATAGCAAGTTAAAATAAGGC. The library was then prepared according to Sugiura *et.al.* (Sugiura et al., 2022). Briefly, the vector was cloned into a pMx-U6-gRNA-BFP retroviral packaging vector and the resulting plasmid library was amplified by electroporated into ElectroMAX DH10B E. coli (Invitrogen 18290015) to maintain greater than 50-fold coverage. Library and screening results are available online at https://functionalimmunogenomics.shinyapps.io/crispr/.

DNA was transfected into Plat-E retroviral packaging cell line using Polyplus jetOPTIMUS DNA and siRNA transfection reagent (Polyplus, 101000051). DMEM media (without puromycin and blasticidin) was changed 4 hours post transfection, and viral supernatant produced by the transfected PlatE cells was collected after an additional 48 hours of culture. CD4^+^ T cells were isolated from the spleen and lymph nodes of OT-II Cas9 double-transgenic mice or Cas9 transgenic mice and activated with splenocytes irradiated at 30Gy and OVA peptide (10mg/mL) (Sigma Aldrich O1641-5mg) for OT-II lung inflammation model. 48 hours post T cell activation, the viral supernatant was spun onto retronectin (Takara Bio, T100B) coated non-tissue culture plates at 2000 x g for 2 hours at 32°C. Activated T cells were transferred to the plates and spun for another 15 minutes at 2000 x g at 32°C, then put in the incubator at 37°C, 5% CO_2_. Either one or two days post T cell transduction, a sample of the cells were collected at 1000-fold representation of the library. Additionally, 3 million live GFP^+^ or BFP^+^ transduced T cells were adoptively transferred into each *Rag1^-/-^*mice by intraperitoneal (i.p.) injection. On days 1, 3, 5, and 7 post-adoptive transfer, mice were sensitized with intranasal ovalbumin protein, resulting in CD4^+^ T cells infiltrating the lung. On day 8 post-adoptive transfer, lungs and spleens were collected, and T cells isolated by positive selection using CD4 (Miltenyi, L3T4) microbeads.

For the IBD model, protocol was followed as above except naïve CD4^+^ T cells were activated with RPMI Th17 skewing media and cytokines and once i.p. injected, mice were weighed twice weekly until 20% weight loss. CD4^+^ T cells from the spleen, mesenteric lymph nodes, and lamina propria were isolated and T cells were isolated using a positive CD4^+^ selection kit.

CRISPR Screen DNA isolation and processing was performed according to Sugiura et.al. (Sugiura et al., 2022). Briefly, genomic DNA from cells were extracted (Millipore Sigma, EXPEXTKB) and gRNA sequences were amplified by two rounds of PCR. Two technical replicates were analyzed with the first amplification round attaching adaptor primers and the second amplification attaching barcoded Illumina sequencing primers. The two amplicons were then purified by gel extraction, combined at equimolar, and sequenced for 150 cycles in paired-end mode on the Illumina Novaseq6000 platform at the Vanderbilt Technologies for Advanced Genomics (VANTAGE) core. At least 1000-fold representation of the library was maintained throughout the process.

Fastq files were analyzed using the Model-based Analysis of Genome-wide CRISPR/Cas9 Knockout (MAGeCKv0.5.0.3) method (Li et al., 2014) to determine statistical significance. The gRNA frequencies in the CD4^+^ T cells isolated from the diseased tissue was compared to the gRNA frequency in the CD4^+^ T cells at the time of adoptive transfer (day 0).

### CRISPR/Cas9 Single Guide Knockout

Single gRNA knockouts were performed as above with the exception that sgRNAs specific to *Mecr* or NTC were used instead of the CRISPR library with the following sequences of sg*Mecr* oligo cloned into a pMx-U6-gRNA-BFP: *Mecr* F CACCGCGTGGCGGTACCAAGCCTCG *Mecr* R AAACCCAGGCTTGGTACCGCCACGC. Five days post-transduction, cells achieve knockout and are used for subsequent experiments.

### T Cell Activation and Differentiation

Primary CD4^+^ T cells were activated using plate-bound anti-CD3 (3mg/mL) (Invitrogen, 16-0031-86) and anti-CD28 (2mg/mL) (Invitrogen, 16-0281-86) antibodies at 1 million cells/well in a tissue culture treated 24-well plate (Thermo Scientific, 142475). 10ng/mL recombinant human IL-2 (Frederick National Laboratory, RO 23-6019), was added to bulk CD4 T cell cultures. Naïve T cells were skewed to the following subsets with cytokines: Th1 (10ng/mL mIL-12 (Biolegend, 577004), 10ng/mL recombinant human IL-2, 1µg/mL anti-IFNγ (Bio X Cell, BP0055), 10µg/mL anti-IL-4 (Bio X Cell, BP0045), Th2 (10ng/mL recombinant human IL-2, 80ng/mL mIL-4 (Peprotech, 214-14), 10µg/mL anti-IFNγ, 10µg/mL anti-IL-12) (Invitrogen, 16-7123-85), Th17 (50ng/mL mIL-6, (Miltenyi, 130-096-685), 10ng/mL mIL-23 (Miltenyi, 130-096-677), 10ng/mL mIL1ß (Miltenyi, 130-094-053), 1ng/mL human TGF-ß1 (Peprotech, 100-21), 10µg/mL anti-IFNγ, 10µg/mL anti-IL-4), iTreg (10ng/mL recombinant human IL-2, 1.5ng/mL human TGF-ß1, 10µg/mL anti-IFNγ, 10µg/mL anti-IL-4).

### Inflammatory Bowel Disease (IBD) Adoptive Transfer

Donor cells were prepared by isolating naïve CD4^+^ T cells (CD4^+^ CD44^-^) by magnetic bead isolation (Miltenyi, 130-104-453) from Cas9 transgenic mice. Cells were polarized to Th17 subset and cultured for 2 days. Cas9-expressing Th17 CD4^+^ T cells were transduced with the PlatE retroviral supernatant containing the CRISPR library or CRISPR sgRNA for *Mecr* or NTC in BFP and cultured for 2 days. sgRNAs for NTC was used in GFP and sg*Mecr* in BFP and 4 million total live cells at an equal amount of transduced T cells each were transferred into *Rag1^-/-^*mice by i.p. injection. Mice were weighed twice weekly to monitor disease progression. Mice were euthanized after 8 weeks, and spleen, mesenteric lymph nodes, lamina propria, intraepithelial lymphocytes, and skin were collected for T cell phenotyping.

### MC-38 OVA Adoptive Transfer Tumor Model

*Rag1^-/-^* were injected subcutaneously with 1 million MC38-OVA tumor cells. After 16 days, CD8^+^ T cells were isolated from the spleen of an OT-I mouse using a negative isolation kit (Miltenyi, 130-095-236) according to manufactures protocol and transduced with the lipid metabolism CRISPR library or sgRNAs in a 1:1 ratio of NTC: *sgMecr*. Mice were injected with 10 million transduced CD8^+^ OT-1 T cells retro-orbitally. Mice were taken down at humane endpoint or after 7 days. Tumors were chopped, mechanically dissociated on a the Miltenyi gentleMACS Octo Dissociator with Heaters (setting implant tumor one) and digested with 435 U/mL DNase I (Sigma-Aldrich, D5025) and 218 U/mL collagenase (Sigma-Aldrich, C2674) at 37°C for 30 min. Tumors were then passed through a 70μm filter and lysed usign ACK lysis buffer. CD8^+^ tumor-infiltrating lymphocytes were isolated using a positive selection kit (Miltenyi, 130 116 478) according to manufactures protocol and CRISPR library preparation and sequencing was followed as above. For 1:1 experiment, purified CD8^+^ T cells were quantified by flow cytometry.

### Listeria-monocytogenes-OVA (Lm-OVA) Adoptive Transfer and Infection

Donor cells were prepared by isolating CD8^+^ T cells by magnetic bead isolation (Miltenyi, 130-104-075) from OT-I;Cas9 transgenic mice. Cells were cultured for 2 days and transduced with a the retroviral supernatant containing a CRISPR sgRNA for NTC in GFP and sg*Mecr* in BFP and cultured for 24 hours. Cells were 1:1 mixed and Rag1^-/-^ mice were i.p. injected with 1 million live transduced cells after mixing. After 5 days in vivo, 5 mice were sacrificed and spleens were isolated. After 30 days total another 5 mice were sacrificed and while the 5 mice were i.p. injected with 1×10^7^ CFU *Listeria monocytogenes* expressing OVA (LM-OVA), as previously described (Madden et al., 2023). The Lm-OVA treated cohort of mice were then sacrificed for analysis 5 days post-infection.

### Flow Cytometry

Single cell suspensions were incubated in Fc block (1:50, BD, 553142) in 50uL for 10 min at room temp, stained for surface markers in 50uL of FACS buffer for 15 min at room temp, washed once with FACS buffer (PBS +2% FBS), and resuspended in 200uL FACS buffer for analysis on a Miltenyi MACSQuant Analyzer 10 or 16. The eBioscience™ Foxp3/transcription factor staining buffer kit (Thermo Fisher, 00-5523-00) was used for intracellular staining of transcription factors. For intracellular cytokine staining, single cell suspensions were incubated for 4 hours at 37°C and 5% CO2 in supplemented RPMI 1640 media supplemented with PMA (50ng/mL, Sigma Aldrich, P8139-1MG), ionomycin (750ng/mL, Sigma Aldrich, I0634-1MG), and GolgiPlug (1:1000, BD Biosciences, 555029) and processed using the BD Cytofix/Cytoperm™ Fixation and Permeabilization Solution (Thermo Fisher, BDB554722). Surface staining was performed as described above. Afterwards, cells were fix/permed for 20 min at 4°C using 50uL of the Cytofixation buffer, and then stained for intracellular markers in 50uL diluted Cytoperm buffer for at least 30 min at 4°C. For flow cytometry on BFP and GFP expressing cells with intracellular transcription factors, cells were fixed in 2% PFA for 20 min room temp before using the FoxP3/transcription factor kit to preserve BFP/GFP signal. Ghost Dye Red 780 viability dye (1:4000, Cell Signaling, 18452S) was used identically to surface antibodies. The antibodies used were: CD4 BV510 (1:200 BD Biociences, 563106), CD4 SB600 (1:200 Invirogen, 2442209), CD4 e450 (1:200 Invitrogen, 48-0041-82), CD8a BV510 (1:200 BD Biociences, 563068), CD8a APC (1:200 BioLegend, 100712), CD62L PE (1: 200 BioLegend, 104408), CD62L APC (1:200 BioLegend, 104412), CD44 PE-Cy7 (1:300 BioLegend, 103030), PD1 PE (1: 200 eBiociences, 12-9985-83), CD71 PE-Cy7 (1:200 BioLegend, 113812), CD25 PE-Cy5 (1:100 Invitrogen, 15-051-82), CD45 FITC (1:200 BioLegend, 157214), CD127 PECy7 (1:50 BioLegend, 135014), KLRG1 APC (1:100 Invitrogen, 17-5893-82), Tbet PE-Cy7 (1:100 BioLegend, 64484), RORγt APC (1:100 Invitrogen, 17-6988-82), FoxP3 PE-Cy5 (1:100 Invitrogen, 15-5773-82), Gata3 PE (1:100 Invitrogen, 1-9966-62), TCF1 AF647 (1:100 Cell Signaling Technologies, C63D9), Eomes PE (1:100 Invitrogen, 12-4875-82), Annexin V APC (1:100 BioLegend, 64090), IFNγ APC (1:100 BD Biociences, 554413), IL-2 PE-Cy5 (1:100 BioLegend, 50384), IL-4 PE-Cy7 (1:100 BioLegend, 504118), IL-17a PE (1:100 Invitrogen, 12-7177-81), TNFα PE-Cy7 (1:100 BioLegend, 506324). P-S6 PE (1:100 Cell Signaling Technologies, 5316S), Propidium Iodide (Sigma, P4864-10mL), BioTracker Far Red Labile 2+ (Millipore, SCT037), Mito-FerroGreen (Dojindo, M489-10). Cell Trace Violet (CTV) (Thermo Fisher, C34557) was used at 1:1000 for cell proliferation assays. Mitochondrial mass was measured with 200nM MitoTracker Green FM (Invitrogen, M7514), mitochondrial membrane potential was measured with 150nM TMRE (Lifetech, T-669), mitochondrial reactive oxygen species was measured with MitoSOX (Invitrogen, M36008), reactive oxygen species was measured with CM-H2DCFDA (Lifetech, C6827), and C11 BODIPY (Thermo Fisher, D3861) was used to measure oxidized lipids. Each staining was incubated for 30min at 37°C 5% CO_2_ in PBS. Flow cytometry data were analyzed using FlowJo v10.7.1. Gating strategies were drawn from singlets, lymphocytes, live cells, and CD4 or CD8^+^ T cells.

### ELISA

Production of IFNγ and IL-10 from T cell culture supernatant was measured using Legend Max ELISA Kit Mouse IFNγ (BioLegend, 430807) and Mouse IL-10 (BioLegend, 431417) according to manufacturer’s protocol. Absorbance was read at 450nm and 570nm and values were normalized to cell number.

### Western Blot

Cells were lysed in Pierce^TM^ RIPA Buffer (Thermo Scientific, 89900) with 1:100 Protease and Phosphatase Inhibitor Cocktail (Thermo Scientific, 78442) for 30 min on ice. Lysates were centrifuged for at 13,000 RPM 15 minutes at 4°C to recover supernatant and quantified for protein concentration by BCA assay (Thermo Scientific, 23225). 70µg of protein was loaded per well for polyacrylamide gel electrophoresis using Mini-PROTEAN Precast Polyacrylamide Gels (Bio-Rad, 4561095). Gel was transferred to a low fluorescence PVDF membrane (Cytiva, 10600023) activated in methanol using 1X Tris-Glycine Transfer Buffer (KD Medical, RGF-3395, with 20% methanol). Membranes were blocked for 1 hour with 5% milk with TBS + 0.5% Tween (TBS-T), before incubation with primary antibody overnight at 4°C. Blots were incubated for 1 hr at room temperature with anti-HRP antibodies and were visualized using Chemiluminescence Pico Plus (Thermo Scientific, 34580) via Amersham Imager 600. The antibodies used for westerns were: ß-actin (1:4000 Cell Signaling Technologies, 3700), Vinculin (1:10,000 Abcam, ab129002), MECR (1:500 Thermo Fisher, 14932-1-AP), OXPHOS cocktail (1:250 Abcam, ab110413), Lipoic Acid (1:1000 Millipore, 437695), Anti-Mouse IgG HRP (1:20,000 Promega W402B), Anti-Rabbit IgG HRP (1:20,000 Promega W401B). Western blot quantification was done by using ImageStudioLite.

### Extracellular Flux Assay

T cells were plated at 150,000 live cells/well in eight technical replicates on a Cell-Tak-coated plate (Corning, 354240) in Agilent Seahorse RPMI 1640 (Agilent, 103576-100) supplemented with 10mM glucose (Agilent, 103577-100), 1mM sodium pyruvate (Agilent, 103578-100), and 2mM glutamine. Cells were analyzed on a Seahorse XFe 96 bioanalyzer using the Mitostress assay (Agilent, 103015–100) with 1μM oligomycin (Cayman Chemical, 11342), 2μM FCCP (Cayman Chemical, 15218), and 0.5μM rotenone (Cayman Chemical 13995)/antimycin A (Cayman Chemical, 19433). Data was then analyzed in the Agilent Wave software version 2.6.

### Targeted Metabolomics

Isolated CD4^+^ T cells from *Cd4*^cre^ – or + *Mecr*^fl/fl^ were plated in culture dishes at 1 million cells per well and cultured for three days with rhIL-2. Cells were then transferred to 15 mL conical tubes, pelleted, and gently washed with room temperature PBS. After washing, cells were pelleted, PBS was removed, samples were flash frozen in liquid N_2_, and stored in a −80°C freezer until ready for metabolite extraction. To extract, 2 mL of - 80°C 80:20 MeOH:H_2_O was added to the cells and samples were vortexed vigorously. 100 nmol ^13^C-1-Lactate was then spiked into each samples as an internal standard (Millipore Sigma, 738778-1G). Cells were then placed in a −80°C freezer to extract for 15 minutes. Precipitated protein from extracted cells was pelleted at 2,800 x g for 10 minutes at 4°C. The supernatant containing extracted metabolites was then transferred to a new 15 mL conical tube containing Extracts were then dried under N_2_. Precipitated protein pellets were resolubilized and protein concentration was measured via BCA assay (Thermo Scientific, 23225). Samples were resuspended in 80 µL 3:2 mobile buffer A: mobile buffer B (see below). 18 µL of the sample was then chromatographed with a Shimadzu LC system equipped with a 100 x 2.1mm, 3.5μm particle diameter XBridge Amide column (Waters, 186004860). Mobile phase A: 20 mM NH_4_OAc, 20 mM NH_4_OH, 5% acetonitrile in H_2_O, pH 9.45; mobile phase B: 100% acetonitrile. With a flow rate of 0.45 mL/min the following gradient was used: 2.0 min, 95% B; 3.0 min, 85% B; 5.0 min, 85% B; 6.0 min, 80% B; 8.0 min, 80% B; 9.0 min, 75% B; 10 min, 75% B; 11 min, 70% B; 12 min, 70% B; 13 min, 50% B; 15 min, 50% B; 16 min 0% B; 17.5 min, 0% B; 18 min, 95% B. The column was equilibrated for 3 minutes at 95% B between each sample. Scheduled MRM was conducted in negative mode with a detection window of 120 seconds using an AB SCIEX 6500 QTRAP with the analyte parameters below. All analytes were quantified via LC-MS/MS using the ^13^C-1-Lactate internal standard and normalized to the protein in each respective sample’s cell pellet. Outliers were removed using interquartile range.

| Species | Q1 (m/z) | Q3 (m/z) | RT (min) |
| --- | --- | --- | --- |
| Glucose | 179 | 89 | 6.9 |
| Hexose Phosphates | 259 | 97 | 13.4 |
| Fructose 1,6-Bisphosphate | 339 | 97 | 13.8 |
| Triose Phosphates | 169 | 79 | 13 |
| 3-Phosphoglycerate | 185 | 97 | 13.4 |
| Phosphoenolpyruvate | 167 | 79 | 13.3 |
| Pyruvate | 87 | 43 | 3.0 |
| Lactate | 89 | 43 | 6.0 |
| <sup>13</sup>C-1-Lactate (Internal Standard) | 90 | 43 | 6.0 |
| Acetyl-CoA | 808 |  |  |
| Citrate/Isocitrate | 191 | 111 | 13.4 |
| cis-Aconitate | 173 | 129 | 12.5 |
| Itaconate | 129 | 85 | 9.7 |
| α-Ketoglutarate | 145 | 101 | 10.3 |
| Succinyl-CoA | 866 |  |  |
| Succinate | 117 | 73 | 10.9 |
| Fumarate | 115 | 71 | 11 |
| Malate | 133 | 115 | 11.4 |
| Oxaloacetate | 131 | 87 | 4.8 |
| Glutamine | 145 | 127 | 9.5 |
| Glutamate | 146 | 102 | 11 |
| Aspartate | 132 | 88 | 11.2 |
| Glucosamine 6-Phosphate | 258 | 97 | 13.4 |
| N-Acetylglucosamine | 220 | 119 | 5.5 |
| N-Acetylglucosamine-Phosphate | 300 | 199 | 12.66 |
| Uridine Diphosphate-N-Acetylglucosamine | 606 | 159 | 12.5 |
| Cyclic AMP | 328 | 134 | 11.2 |
| AMP | 346 | 79 | 12 |
| ADP | 426 | 328 | 13.3 |
| ATP | 506 | 408 | 13.6 |
| NAD<sup>+</sup> | 663 | 540 | 12.3 |
| NADP<sup>+</sup>/NADPH | 742 | 620 | 13.6 |
| CoASH | 766 |  | 13 |
| GTP | 522 | 79 | 13.7 |
| GDP | 422 | 79 | 12.8 |
| FAD | 784 | 437 | 11.4 |
| GSH | 306 | 128 | 11.1 |
| GSSG | 611 | 306 | 13.7 |

### Ferroptosis Rescue

CD4^+^ T cells from *Mecr*^fl/fl^; *Cd4*^cre^ mice were isolated and activated with anti-CD3, anti-CD28, and rhIL-2 in complete RPMI 1640 media. Cells were cultured for 3 days and treated with mouse anti-CD71 blocking antibody or isotype control as previously described (Voss et al., 2023). Cells were then washed in PBS and resuspended at 1×10^6^ cells/mL in fresh media and pre-treated for 2 hours with 0.5mM Glutathione ethyl ester (Et-GSH, Cayman Chemical, 14953) or 50nM UAMC-3203 (Cayman Chemical, 26525) to prevent ferroptosis. 0.5µM (1S,3R)-RSL3 (Cayman Chemical, 19288) or DMSO solvent control was then applied overnight to induce ferroptosis. Viability and markers of ferroptosis were quantified the next day by flow cytometry.

### Quantification and Statistical Analysis

Graphs and statistical tests were generated using GraphPad Prism 10 unless otherwise noted. Sample sizes were chosen based on previous studies. Statistical significance was performed by unpaired two-sided t tests, two-way ANOVA with Tukey post-hoc test, Dunnett’s post-hoc test, or Welch’s t test. Graphs show mean and SEM unless otherwise stated. ns = p value >0.05, * = p value <0.05, ** = p value <0.01, *** = p value <0.001, **** = p value <0.0001.

### Data Availability

Custom gRNA library information and resulting CRISPR/Cas9 screening data are available on https://functionalimmunogenomics.shinyapps.io/crispr/. Data used for analysis in Supplemental Figure 1 was from the Immunological Genome Project database (ImmGen) (Heng et al., 2008) (a), GSE232241 (Madden et al., 2023) (b), GSE41870 (Doering et al., 2012) (c), Immunological Proteomic Resourse (ImmPRes) (Brenes et al., 2023) (d, e).

## Acknowledgements

We thank the members of the J. Rathmell lab and Dr W. Kimryn Rathmell for contributing to this project and input. We also thank Drs Russell Jones and Sarah Nowinski (Van Andel Institute) for their suggestions and input. For assistance with CRISPR/Cas9 screen sequencing, we acknowledge the Vanderbilt Technologies for Advanced Genomics (VANTAGE). We acknowledge the Translational Pathology Shared Resource, which is supported by NCI/NIH Cancer Center Support Grant P30CA068485. Diagrams were created using Biorender.

## Disclosures

Dr. J.C. Rathmell is a founder and scientific advisory board member of Sitryx Therapeutics.

## Supplemental Figure Legends

**Supplemental Table 1: CRISPR/Cas9 Lipid metabolism gRNA library.** Table shows list of all genes and positive controls in the library as well as the relevent corresponding lipid metabolism pathway.

**Supplemental Figure 1:**
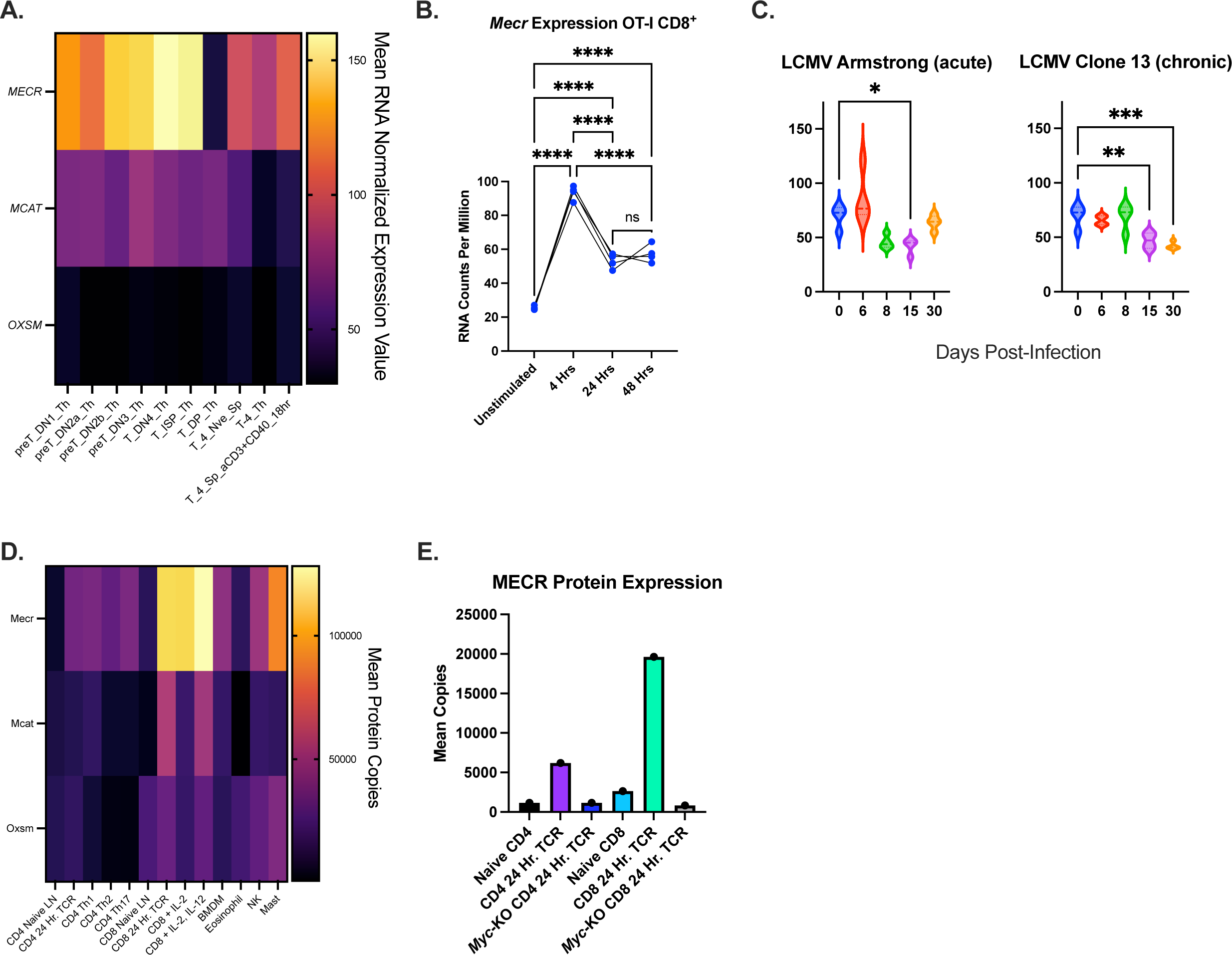
*Mecr* expression changes upon T cell activation. **a)** Mean normalized expression values of mtFAS human genes from the Immunological Genome Project database (ImmGen) (Heng et al., 2008). Labels are the following: preT_DN1_Th-Thymic preT Double Negative 1, preT_DN2a_Th-Thymic preT Double negative 2a, preT_DN2b_Th-Thymic preT Double Negative 2b, PreT_DN3_Th-Thymic preT Double Negative 3, T_DN4_Th-Thymic Double Negative 4, T_ISP-Th-Intermediate single positive thymocytes, T_DP_Th-Thymic double positive, T_4_Nve_Sp-Splenic Naïve CD4, T-4_Th-CD4+ Thymocytes, T_4_Sp aCD3_CD40_18hr-Splenic activated CD4^+^ T cells, aCD3+CD40 18 Hr. **b)** Quantified *Mecr* expression time course in activated antigen-specific murine OT-1 CD8^+^ T cell RNAseq from GSE232241 (Madden et al., 2023). **c)** Expression levels of *Mecr* mRNA in CD8^+^ T cells that specifically recognize H2-Db GP33 LCMV antigen during infection. CD8^+^ T cells were isolated from mice infected with acute (LCMV-Arm) or chronic (LCMV-C13) variants of the virus at different days post infection. The data are reanalyzed from GSE41870 (Doering et al., 2012) **d)** Mean human protein copies of mtFAS genes from the Immunological Proteomic Resourse (ImmPRes) (Brenes et al., 2023). **e)** Quantified MECR expression in murine T cells with or without myc-KO from ImmPRes. Panel (b) each data point represents a biological replicate and error bars show standard deviation. Panel (b) statistical significance performed by one-way ANOVA with Tukey’s multiple comparison post-hoc test. Panel (c) statistical significance performed by one-way ANOVA with Dunnett’s multiple comparison post-hoc test. (* p<0.05, ** p<0.01, *** p<0.001, **** p<0.0001).

**Supplemental Figure 2:**
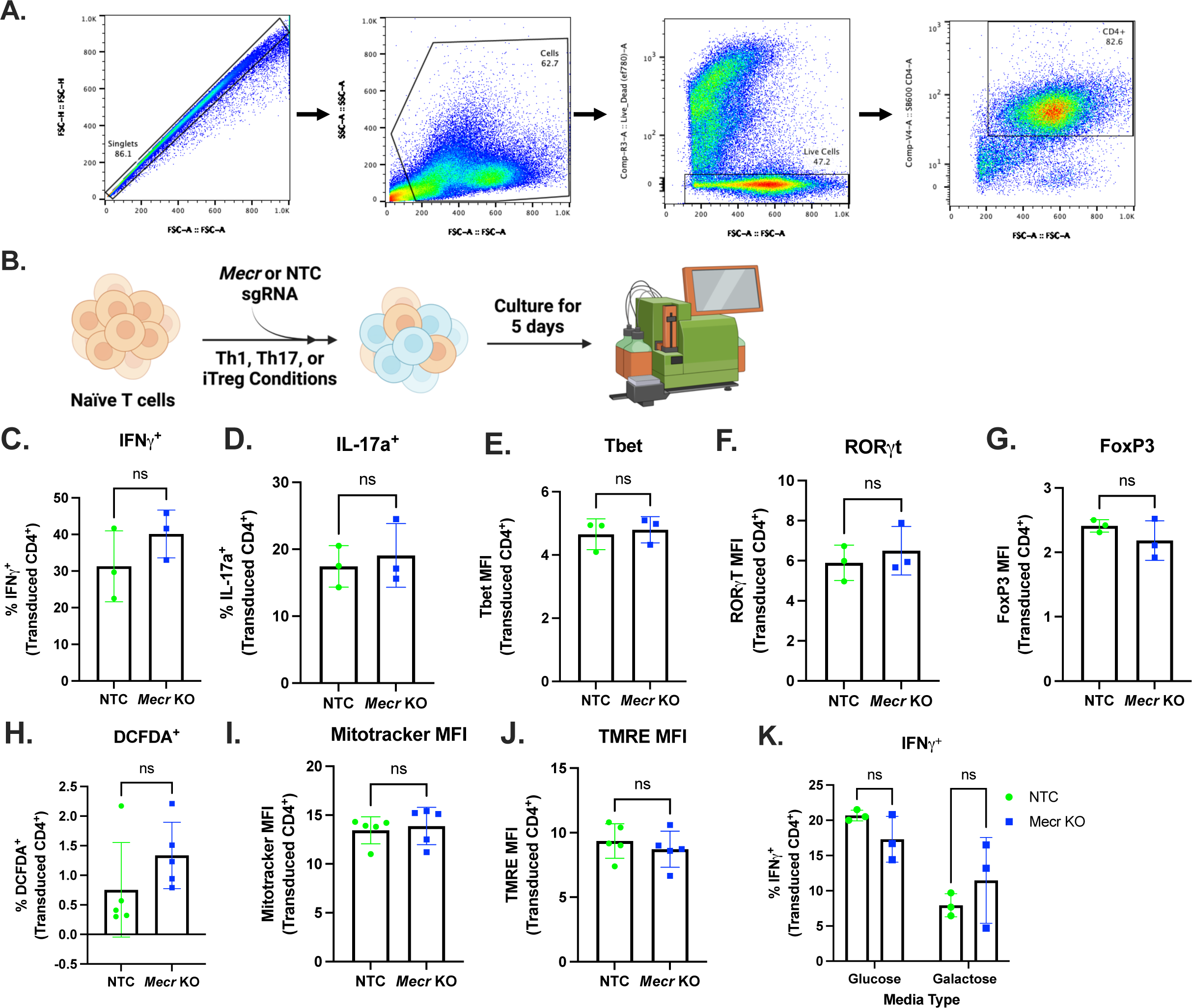
*In vitro* disruption of *Mecr* with CRISPR/Cas9 does not alter T cell function. **a)** Gating scheme for all experiments. **b)** Experimental scheme of NTC or *Mecr*-KO *in vitro* CRISPR/Cas9 protocol. **c-g)** Differentiated naïve CD4^+^ T cells transduced with sgNTC or sg*Mecr*. **c)** Percentage of IFNγ^+^ Th1. **d)** Percentage of IL-17a^+^ Th17. **e)** Tbet MFI of Th1. **f)** RORγt MFI Th17. **g)** FoxP3 MFI iTreg. **h-j)** Mitochondrial stains of activated CD4^+^ T cells transduced with sgNTC or sg*Mecr*. **h)** Percentage of DCFDA^+^. **i)** Mitotracker MFI. **j)** TMRE MFI. **k)** Percentage of IFNγ^+^ T cells from CD4^+^ T cells activated and cultured in media with glucose or galactose. Panels (c-k) show representative results from three independent experiments. Each data point represents a biological replicate and error bars show standard deviation. Panels (c-j) statistical significance performed by unpaired t tests. Panel (k) statistical significance performed by two-way ANOVA with Sidak’s multiple comparison. (* p<0.05, ** p<0.01, *** p<0.001, **** p<0.0001).

**Supplemental Figure 3:**
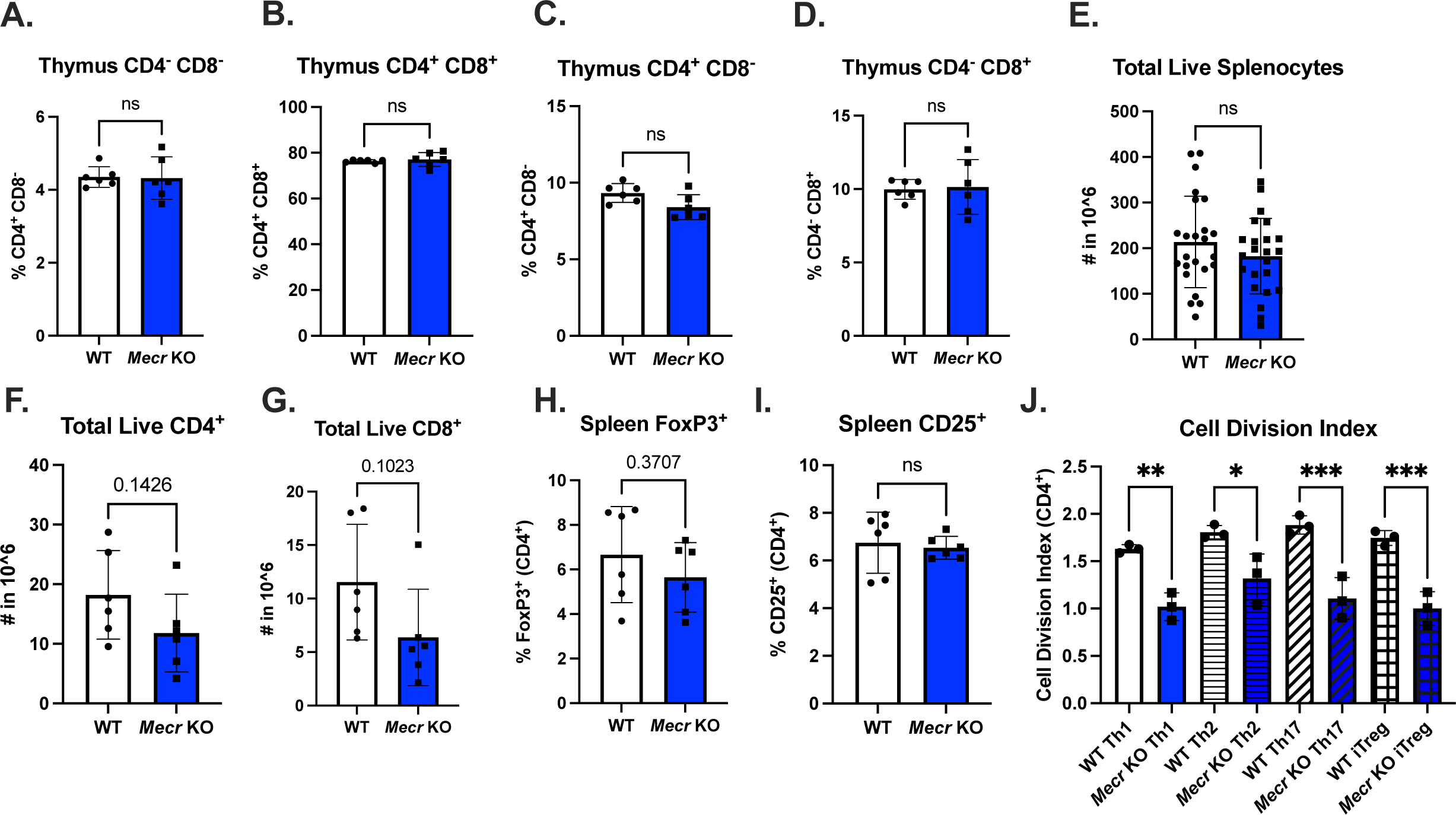
*Mecr*^fl/fl^; *Cd4*^cre^ mice display no difference in thymocytes and T regulatory cells *ex vivo*. **a-d)** CD4/CD8 thymocytes of live cells. **e)** Total live splenocytes of *Mecr*^fl/fl^; *Cd4*^cre^ mice**. f)** Number of live CD4^+^ T cells. **g)** Number of live CD8^+^ T cells. **h, i)** Quantification of T regulatory cells ex vivo of FoxP3^+^ CD4^+^ T cells and CD25^+^ T cells. **j)** Cell division index from CTV in T cell subsets. Panels (a-i) show results from two pooled independent experiments. Panel (j) shows results from one independent experiment. Each data point represents a biological replicate and error bars show standard deviation. Panels (a-i) statistical analysis significance performed by unpaired t tests.Panel (j) statistical analysis for significance performed by two-way ANOVA with Tukey post-hoc test (* p<0.05, ** p<0.01, *** p<0.001, **** p<0.0001).

**Supplemental Figure 4:**
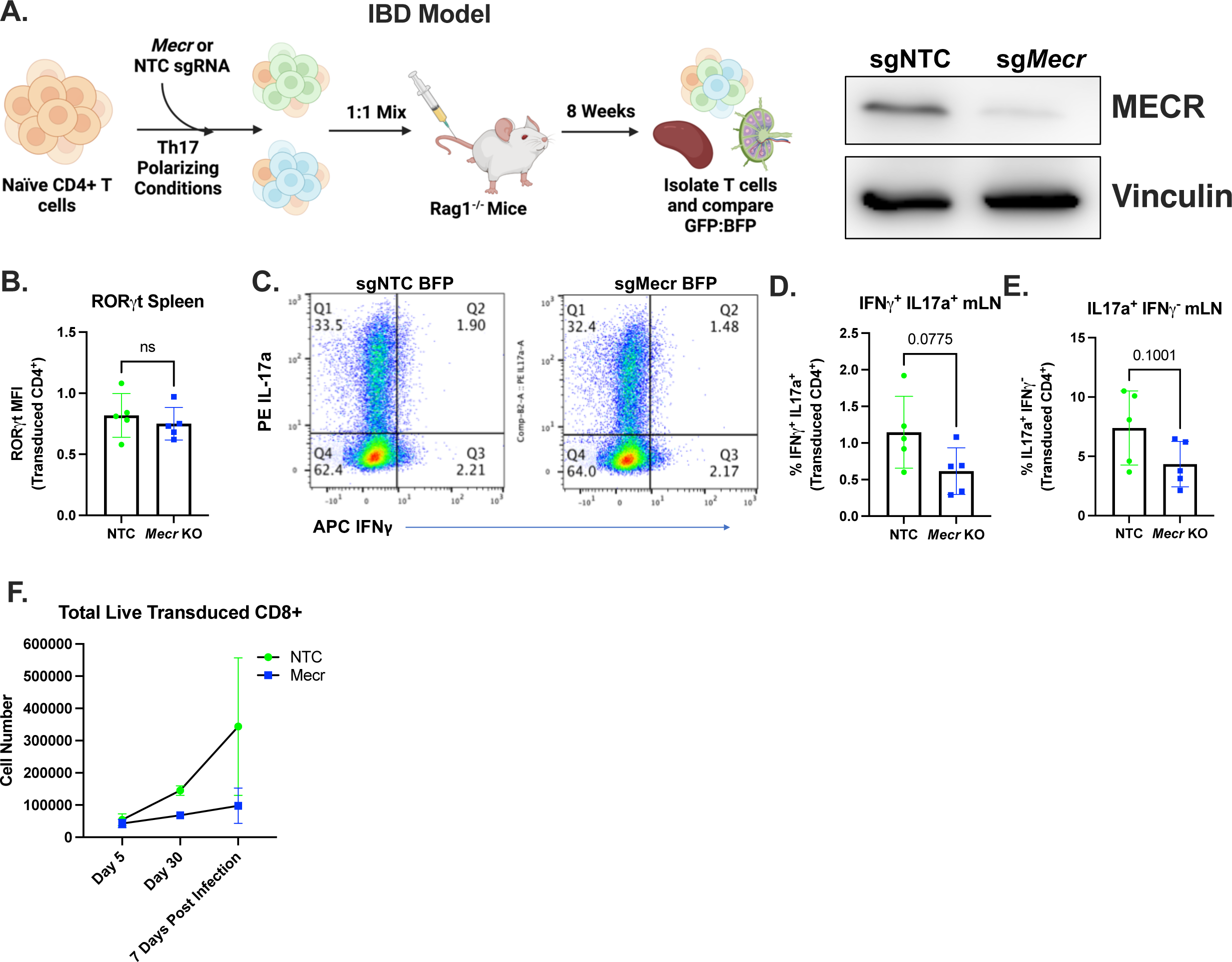
*Mecr*-KO does not cause reduced Th17 and CD8 function in CRISPR/Cas9 1:1 Experiments. **a)** Experimental scheme of *in vivo* 1:1 IBD CRISPR/Cas9 protocol. **b-e)** IBD model. **b)** RORγt expression post-disease. **c)** IL-17a and IFNγ production in skewed Th17 CD4^+^ T cells day of adoptive transfer. **d)** Percentage of IFNγ^+^ IL-17a^+^ post-disease. **e)** Percentage of IL17a^+^ IFNγ^-^ post-disease. **f)** Listeria-OVA CD8^+^ 1:1 total live transduced CD8^+^ T cells. Panels (b-e) show representative results from three independent experiments. Panel (f) shows representative results from two independent experiments. Each data point represents a biological replicate and error bars show standard deviation. Statistical significance performed by unpaired t tests. (* p<0.05, ** p<0.01, *** p<0.001, **** p<0.0001).

## Notes

Funded by R01AI153167, R01DK105550, R01CA217987, R01HL136664, a Mark Foundation Endeavor Award, and the Distinguished Innovator of the Lupus Research Alliance to (JCR), the Juvenile Diabetes Research Foundation Advanced Postdoctoral Fellowship (K.V.), K00CA253718 (E.N.A.), T32CA009582 (EQJ), F30CA239367 (M.Z.M.), T32GM007347 (M.Z.M., A.S.), R38HL143619 (A.C.Y.), and 2T32CA009592-32 (K.K.S.).

