## Supplementary figures and images for "Mitochondrial Fatty Acid Synthesis and Mecr Regulate CD4^+^ T Cell Function and Oxidative Metabolism"

### Supplemental Figures

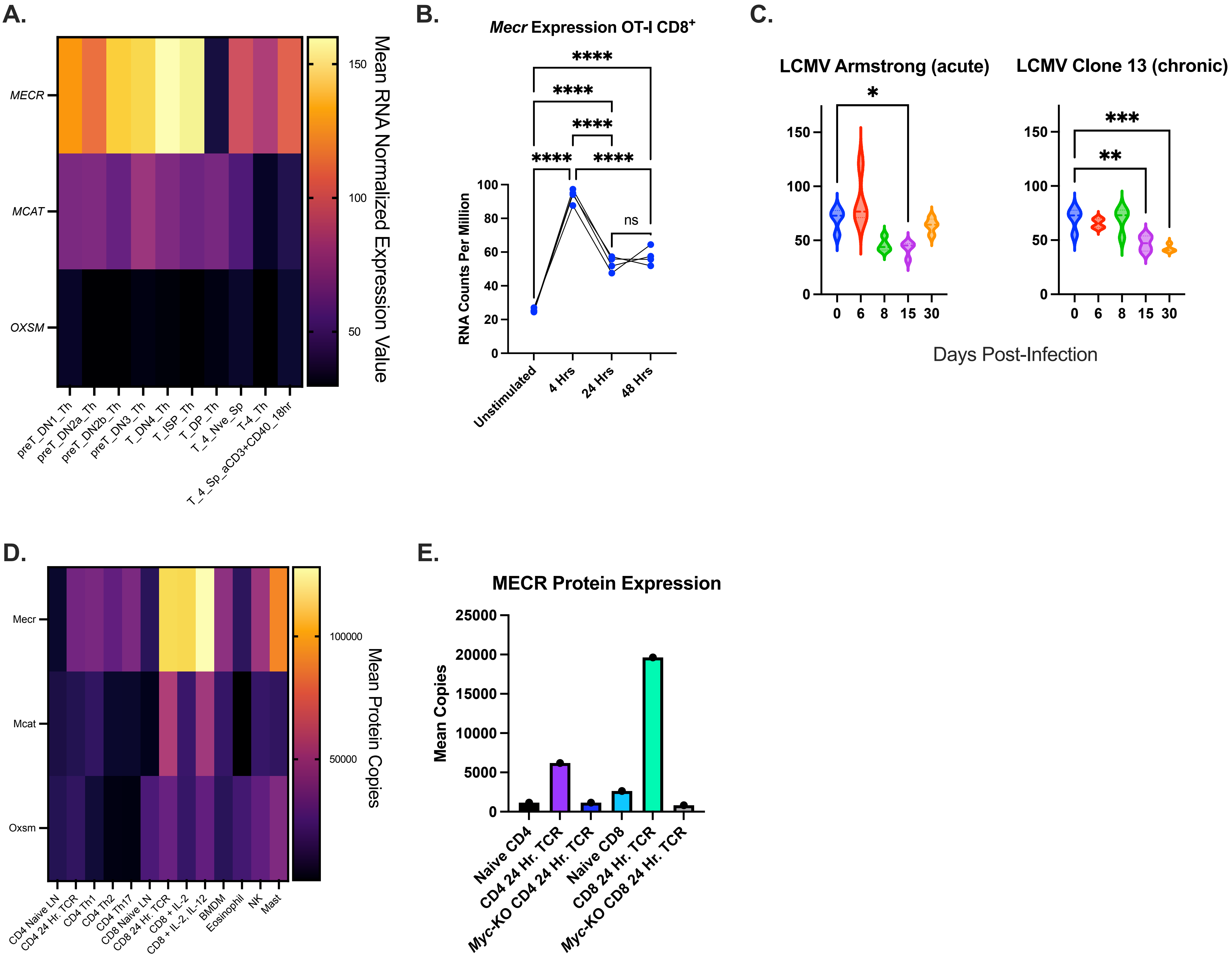

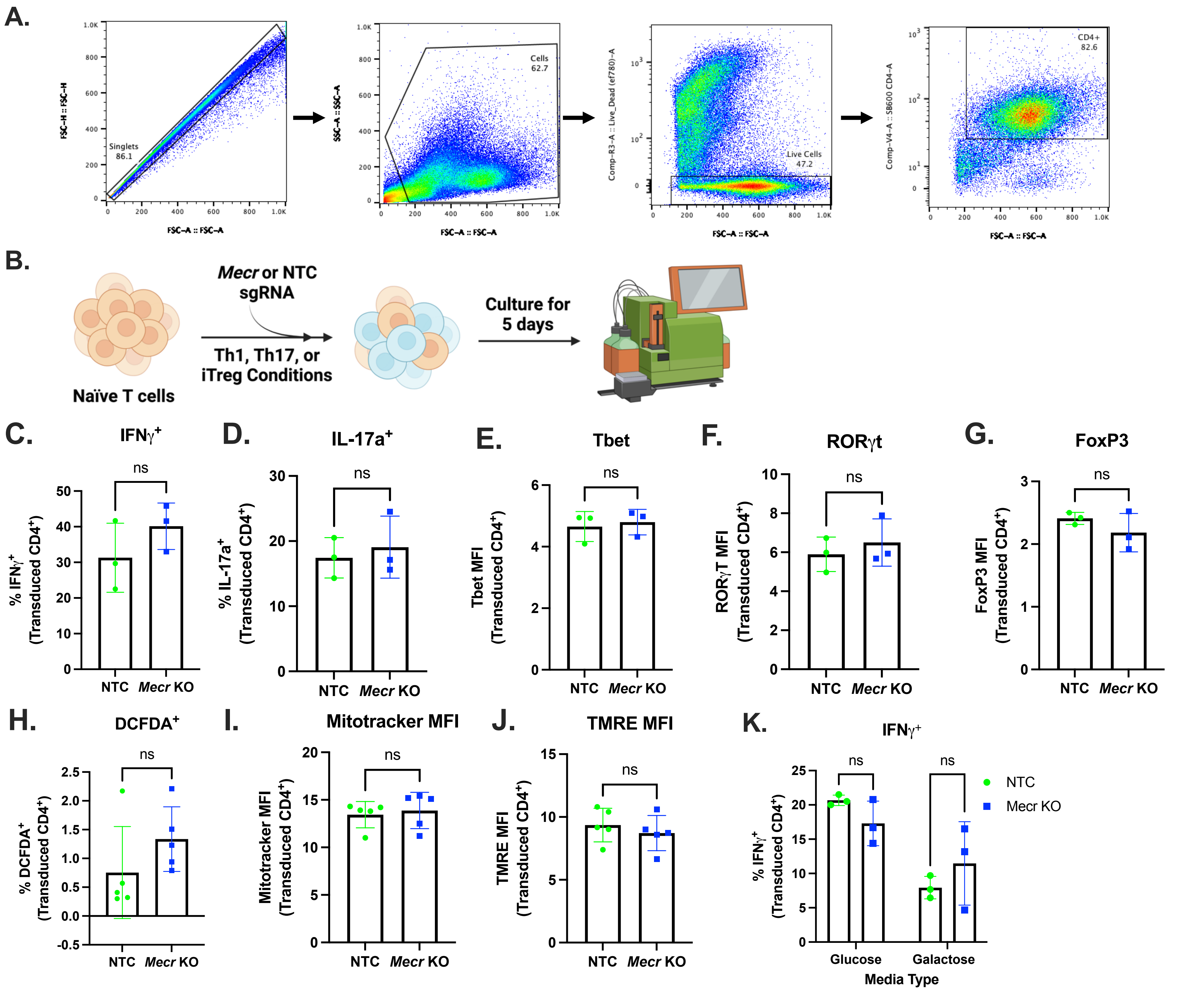

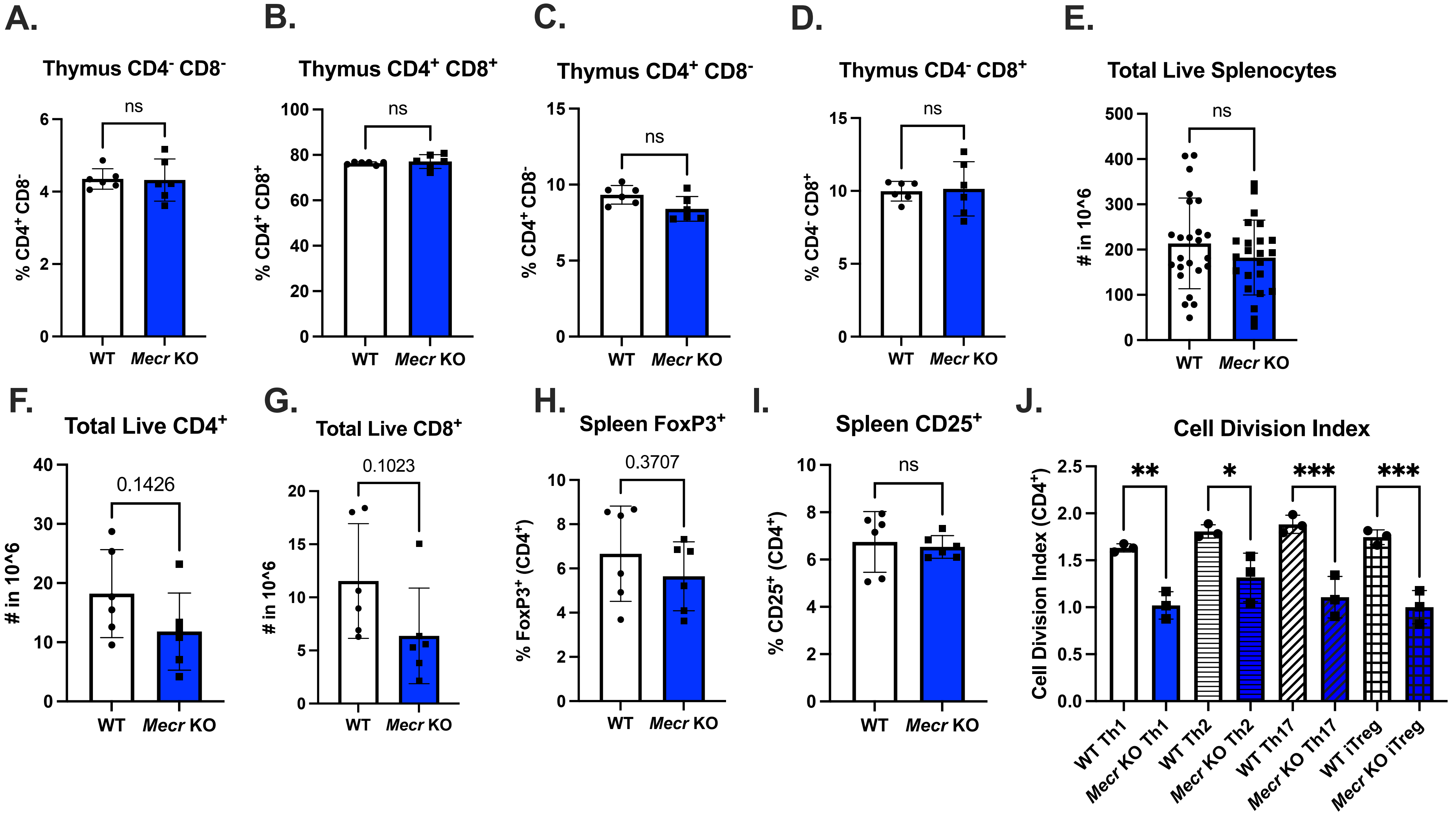

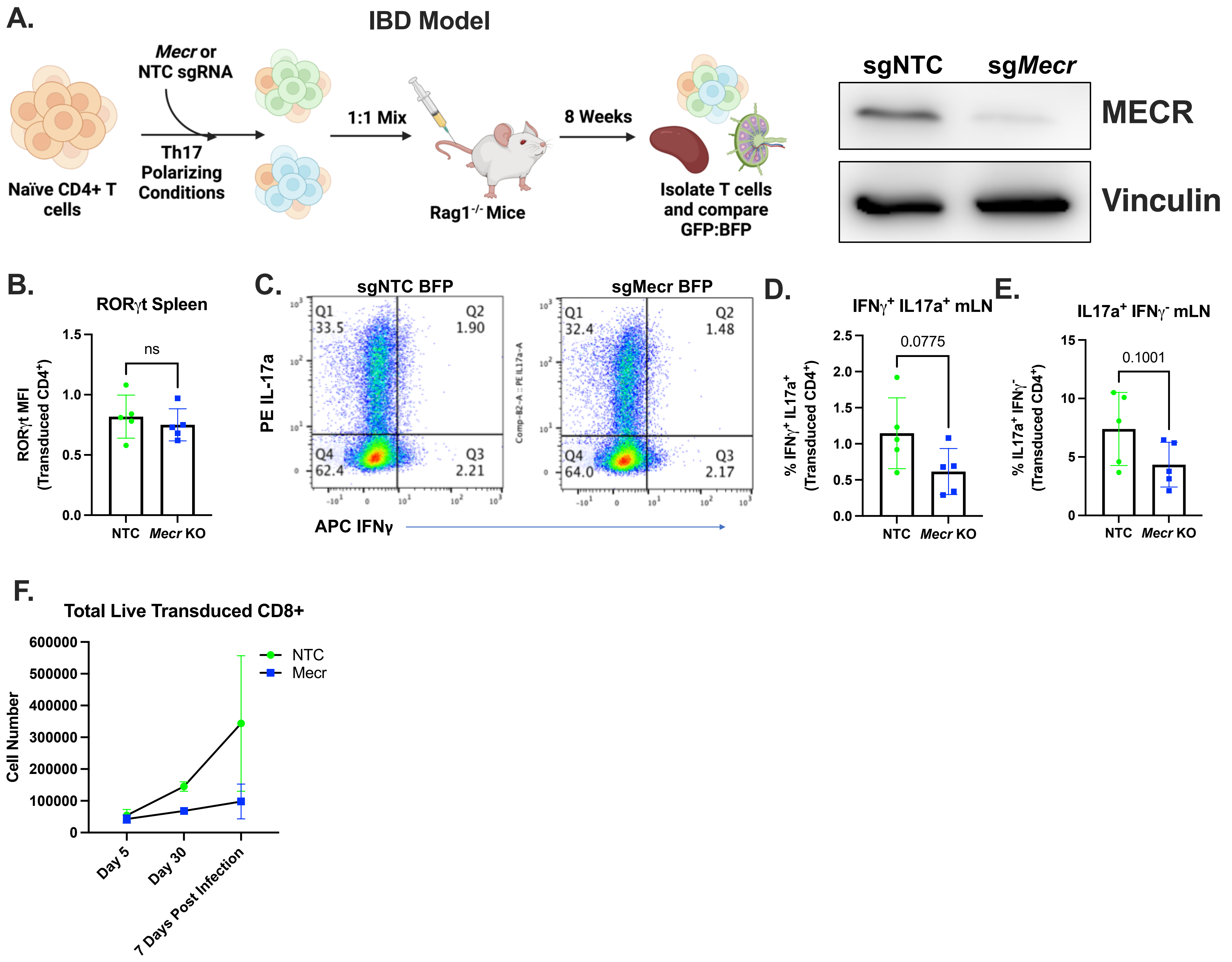
